# DateLife: leveraging databases and analytical tools to reveal the dated Tree of Life

**DOI:** 10.1101/782094

**Authors:** Luna L. Sánchez Reyes, Emily Jane McTavish, Brian O’Meara

## Abstract

Achieving a high-quality reconstruction of a phylogenetic tree with branch lengths proportional to absolute time (chronogram) is a difficult and time-consuming task. But the increased availability of fossil and molecular data, and time-efficient analytical techniques has resulted in many recent publications of large chronograms for a large number and wide diversity of organisms. Knowledge of the evolutionary time frame of organisms is key for research in the natural sciences. It also represent valuable information for education, science communication, and policy decisions. When chronograms are shared in public and open databases, this wealth of expertly-curated and peer-reviewed data on evolutionary timeframe is exposed in a programatic and reusable way, as intensive and localized efforts have improved data sharing practices, as well as incentivizited open science in biology. Here we present DateLife, a service implemented as an R package and an R Shiny website application available at www.datelife.org, that provides functionalities for efficient and easy finding, summary, reuse, and reanalysis of expert, peer-reviewed, public data on time frame of evolution. The main DateLife workflow constructs a chronogram for any given combination of taxon names by searching a local chronogram database constructed and curated from the Open Tree of Life Phylesystem phylogenetic database, which incorporates phylogenetic data from the TreeBASE database as well. We implement and test methods for summarizing time data from multiple source chronograms using supertree and congruification algorithms, and using age data extracted from source chronograms as secondary calibration points to add branch lengths proportional to absolute time to a tree topology. DateLife will be useful to increase awareness of the existing variation in alternative hypothesis of evolutionary time for the same organisms, and can foster exploration of the effect of alternative evolutionary timing hypotheses on the results of downstream analyses, providing a framework for a more informed interpretation of evolutionary results.

## Introduction

Chronograms –phylogenies with branch lengths proportional to time– provide key data on evolutionary time frame for the study of natural processes in many areas of biological research, such as developmental biology (Delsuc et al., 2018; Laubichler & Maienschein, 2009), conservation biology (Felsenstein, 1985; Webb, 2000), historical biogeography (Posadas, Crisci, & Katinas, 2006), and species diversification (Magallon & Sanderson, 2001; Morlon, 2014).

Building a chronogram is not an easy task. It requires obtaining and curating data to construct a phylogeny, selecting and placing appropriate calibrations on the phylogeny using independent age data points from the fossil record or other dated events, and inferring the full dated tree; it also generally requires specialized biological training, taxonomic domain knowledge, and a non-negligible amount of research time, computational resources and funding.

Here we present the DateLife project which has the main goal of capturing age data from published chronograms, and making these data readily accessible to the community for reuse and reanalysis, for research, teaching, and science communication and policy. DateLife’s core software application is available as an R package (Sanchez-Reyes et al., 2022), and as an online Rshiny interactive website at www.datelife.org. It features key elements for scientific reproducibility, such as a versioned, open and fully public source database (McTavish et al., 2015), data stored and available in a computer readable format (Vos et al., 2012), automated and programmatic ways of accessing the data (Stoltzfus et al., 2013) and methods to summarize and compare the data.

## Description

DateLife’s core software application consists of the R package datelife. Its current stable version – v0.6.5, is available from the The Comprehensive R Archive Network (CRAN) repository (Sanchez-Reyes et al., 2022), and relies on functionalities from various biological R packages: ape (Paradis, Claude, & Strimmer, 2004), bold (Chamberlain et al., 2019), geiger (Pennell et al., 2014), paleotree (Bapst, 2012), phyloch (Heibl, 2008), phylocomr (Ooms & Chamberlain, 2018), phytools (Revell, 2012), rotl (Michonneau, Brown, & Winter, 2016), and taxize (Chamberlain & Szöcs, 2013; Chamberlain et al., 2019). Figure 1 provides a graphical summary of the three main steps of the DateLife workflow: creating a search query, searching a database, and summarizing results from the search.

**Figure 1.**
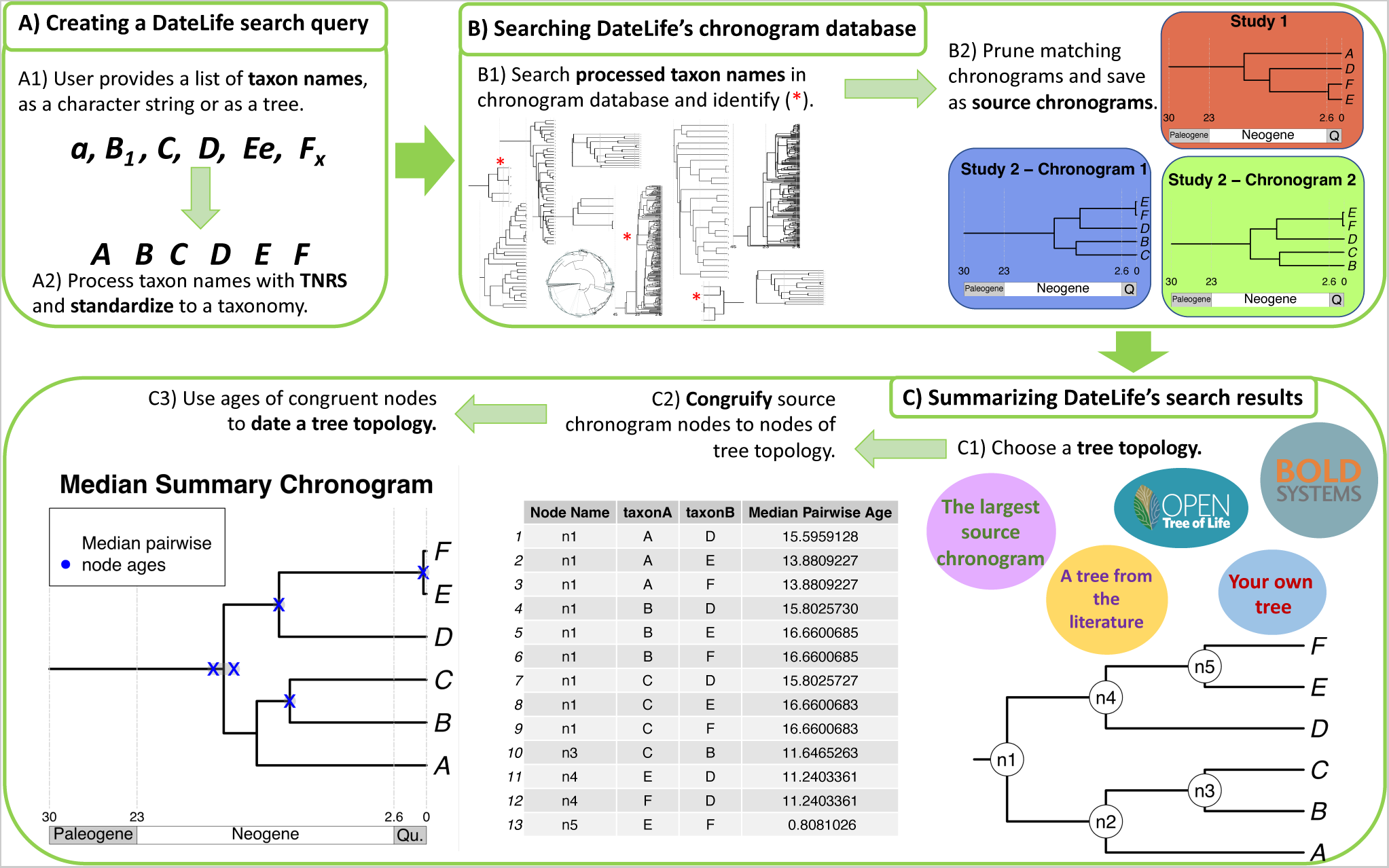
Main DateLife workflow. Analyses can be performed via DateLife’s interactive website at www.datelife.org, or using the `datelife` R package. Details on the R functions used to perform the analyses are available from `datelife`’s R package vignettes at phylotastic.org/datelife.

### Creating a search query

DateLife starts by processing an input consisting of at least two taxon names, which can be provided as a comma separated character string or as tip labels on a tree. If the input is a tree, it can be provided as a classic newick character string (Archie et al., 1986), or as a “phylo” R object (Paradis et al., 2004). The input tree is not required to have branch lengths, and its topology is used in the summary steps described in the next section.

DateLife accepts scientific names that can belong to any inclusive taxonomic group (e.g., genus, family, tribe, etc.) or a binomial species name. Subspecies and variants are ignored. If an input taxon name belongs to an inclusive taxonomic group, DateLife has two alternative behaviors defined by the “get species from taxon” flag. If the flag is active, DateLife retrieves all species names within the inclusive taxonomic group following a standard taxonomy of choice, and adds them to the input string. Taxonomies currently supported by DateLife are Open Tree of Life (OpenTree) unified Taxonomy (OTT, Rees & Cranston, 2017), the National Center of Biotechnology Information (NCBI) taxonomic database (Schoch et al., 2020), the Global Biodiversity Information Facility (GBIF) taxonomic backbone (GBIF Secretariat, 2022), and the Interim Register of Marine and Nonmarine Genera (IRMNG) database (Rees, Vandepitte, Decock, & Vanhoorne, 2017). If the flag is inactive, DateLife excludes any taxon names above the species level from the search query.

DateLife processes input scientific names using a Taxonomic Name Resolution Service (TNRS), which increases the probability of correctly finding the queried taxon names in the chronogram database. TNRS detects, corrects and standardizes name misspellings and typos, variant spellings and authorities, and nomenclatural synonyms to a single taxonomic standard (Boyle et al., 2013). DateLife implements TNRS with OTT as standard (Open Tree Of Life et al., 2016; Rees & Cranston, 2017), storing taxonomic identification numbers for further processing.

The processed input taxon names are saved as an R object of a newly defined class, `datelifeQuery`, that is used in the following steps. This object contains the standardized names, the corresponding OTT identification numbers, and the topology of the input tree if any was provided.

### Searching a chronogram database

At the time of writing of this manuscript (Jun 22, 2022), DateLife’s chronogram database latest version consist of 253 chronograms published in 187 different studies. It is curated from OpenTree’s phylogenetic database, the Phylesystem, which constitutes an open source of expert and peer-reviewed phylogenetic knowledge with rich metadata (McTavish et al., 2015), which allows automatic and reproducible assembly of our chronogram database. Datelife’s chronogram database is navigable as an R data object within the `datelife` R package.

A unique feature of the Phylesystem is that any user can add new published, state-of-the-art chronograms any time, through their curator application (https://tree.opentreeoflife.org/curator). As chronograms are added to Phylesystem, they are incorporated into the chronogram database of the `datelife` package. The updated database is assigned a new version number, followed by a package release on CRAN. `datelife`’s chronogram database is updated as new chronogram data is added to Phylesystem, at a minimum of once a month and a maximum of every 6 months. Users can also implement functions from the `datelife` R package to trigger an update of the local chronogram database, to incorporate any new chronograms to the user’s DateLife analysis before an official database update is released on CRAN.

A DateLife search is implemented by matching processed taxon names provided by the user to tip labels in the chronogram database. Chronograms with at least two matching taxon names on their tip labels are identified and pruned down to preserve only the matched taxa. These matching pruned chronograms are referred to as source chronograms. Total distance (in units of millions of years) between taxon pairs within each source chronogram are stored as a patristic distance matrix (Figure 1). The matrix format speeds up extraction of pairwise taxon ages of any queried taxa, as opposed to searching the ancestor node of a pair of taxa in a “phylo” object or newick string. Finally, the patristic matrices are associated to the study citation where the original chronogram was published, and stored as an R object of the newly defined class `datelifeResult`.

### Summarizing search results

Summary information is extracted from the `datelifeResult` object to inform decisions for subsequent steps in the analysis workflow. Basic summary information available to the user is:

1. The matching pruned chronograms as newick strings or “phylo” objects.
2. The ages of the root of all source chronograms. These ages can correspond to the age of the most recent common ancestor (mrca) of the user’s group of interest if the source chronograms have all taxa belonging to the group. If not, the root corresponds to the mrca of a subgroup withing the group of interest.
3. Study citations where original chronograms were published.
4. A report of input taxon names matches across source chronograms.
5. The source chronogram(s) with the most input taxon names.
6. Various single summary chronograms resulting from summarizing age data, generated using the methodology described next.

#### Choosing a topology

DateLife requires a tree topology to summarize age data upon. We recommend that users provide a tree topology as input from the literature, or one of their own making. If no topology is provided, DateLife automatically extracts one from the OpenTree synthetic tree, a phylogeny encompassing 2.3 million taxa across all life, assembled from 1, 239 published phylogenetic trees and OpenTree’s unified Taxonomy, OTT (Open Tree Of Life et al., 2019). Alternatively, DateLife can combine topologies from source chronograms using a supertree approach. To combine topologies from source chronograms into a single summary (or supertree) topology, the DateLife workflow identifies the source chronograms that form a grove, roughly, a sufficiently overlapping set of taxa between trees, by implementing definition 2.8 for n-overlap from Ané et al. (2009). In rare cases, a group of trees can have multiple groves. By default, DateLife chooses the grove with the most taxa, however, the “criterion = trees” flag allows the user to choose the grove with the most trees instead. If source chronograms do not form a grove, the supertree reconstruction will fail.

#### Dating the topology

Input topologies from OpenTree or the supertree approach described above do not include branch length estimates of any kind. Optionally, to estimate branch lengths proportional to substitution rates on these topologies, DateLife can mine the Barcode of Life Data System, BOLD (Ratnasingham & Hebert, 2007) to obtain genetic markers for the input taxa. These markers are aligned with MUSCLE (Edgar, 2004) (by default) or MAFFT (Katoh, Asimenos, & Toh, 2009). This alignment can be used to estimate branch lengths on input topologies that lack branch lengths. Currently, branch length reconstruction in DateLife is performed using parsimony and the likelihood of the phylogenetic tree given a sequence alignment is computed (Schliep, 2011). While relative branch length information provides additional data for nodes without secondary date calibrations, topologies without branch lengths can also be dated.

Once a topology is chosen, DateLife applies the congruification method (Eastman, Harmon, & Tank, 2013) to find nodes belonging to the same clade across source chronograms, and extract the corresponding node ages from the patristic distance matrices stored as `datelifeResult`. By definition, the matrices store total distance (time from tip to tip), hence, node ages correspond to half the values stored in the patristic distance matrices. This assumes that the terminal taxa are coeval and occur at the present. A table of congruified node ages that can be used as calibrations for a dating analysis is stored as a `congruifiedCalibrations` object.

For each congruent node, the pairwise distances that traverse that node are summarized into a single summary matrix using classic summary statistics (i.e., mean, median, minimum and maximum ages), and the Supermatrix Distance Method [SDM; Criscuolo, Berry, Douzery, and Gascuel (2006)], which deforms patristic distance matrices by minimizing variance and then averaging them. These single summary taxon pair age matrices (Summarized calibrations) can be applied as calibrations to date a tree topology, using different dating methods currently supported within DateLife: MrBayes (Huelsenbeck & Ronquist, 2001; Ronquist & Huelsenbeck, 2003), PATHd8 (Britton, Anderson, Jacquet, Lundqvist, & Bremer, 2007), BLADJ (Webb, Ackerly, & Kembel, 2008; Webb & Donoghue, 2005), and treePL (Smith & O’Meara, 2012).

By default, DateLife implements the Branch Length Adjuster (BLADJ) algorithm to obtain a fully dated topology. BLADJ fixes node ages that have calibration data, and distributes time between nodes with no data evenly between nodes with calibration data. This minimizes age variance in the resulting chronogram (Webb et al., 2008). BLADJ does not use branch lengths even when they are present in the input tree or summarizing topology. When there is conflict in ages between nodes with calibration data, BLADJ ignores node ages that are older than the age of a parent node. BLADJ requires a root age estimate. If there is no information on the age of the root in the chronogram database, users can provide an estimate from the literature. If none is provided, DateLife assigns an arbitrary age to the root as 10% older than the oldest age available within the group.

Alternative phylogenetic dating options supported in DateLife (MrBayes, PATHD8, TreePL) incorporate branch length information from the input topology in combination with the calibrations. PATHd8 is a non-clock, rate-smoothing method (Britton et al., 2007) to date trees. treePL (Smith & O’Meara, 2012), is a semi-parametric, rate-smoothing, penalized likelihood dating method (Sanderson, 2002). The MrBayes (Huelsenbeck & Ronquist, 2001; Ronquist & Huelsenbeck, 2003) approach in DateLife uses the calibrations as priors on node ages.

#### Visualizing results

Finally, users can save all source and summary chronograms in formats that permit reuse and reanalyses (newick and R “phylo” format), as well as visualize and compare results graphically, or construct their own graphs using DateLife’s chronogram plot generation functions available from the R package `datelifeplot` (Sanchez-Reyes & O’Meara, 2022).

## Benchmark

`datelife`’s R package code speed was tested on an Apple iMac with one 3.4 GHz Intel Core i5 processor. We registered variation in computing time of query processing and search through the database relative to number of queried taxon names. Query processing time increases roughly linearly with number of input taxon names, and increases considerably if Taxonomic Name Resolution Service (TNRS) is activated. Up to ten thousand names can be processed and searched in less than 30 minutes with the most time consuming settings. Once names have been processed as described in methods, a name search through the chronogram database can be performed in less than a minute, even with a very large number of taxon names (Fig. 2).

**Figure 2.**
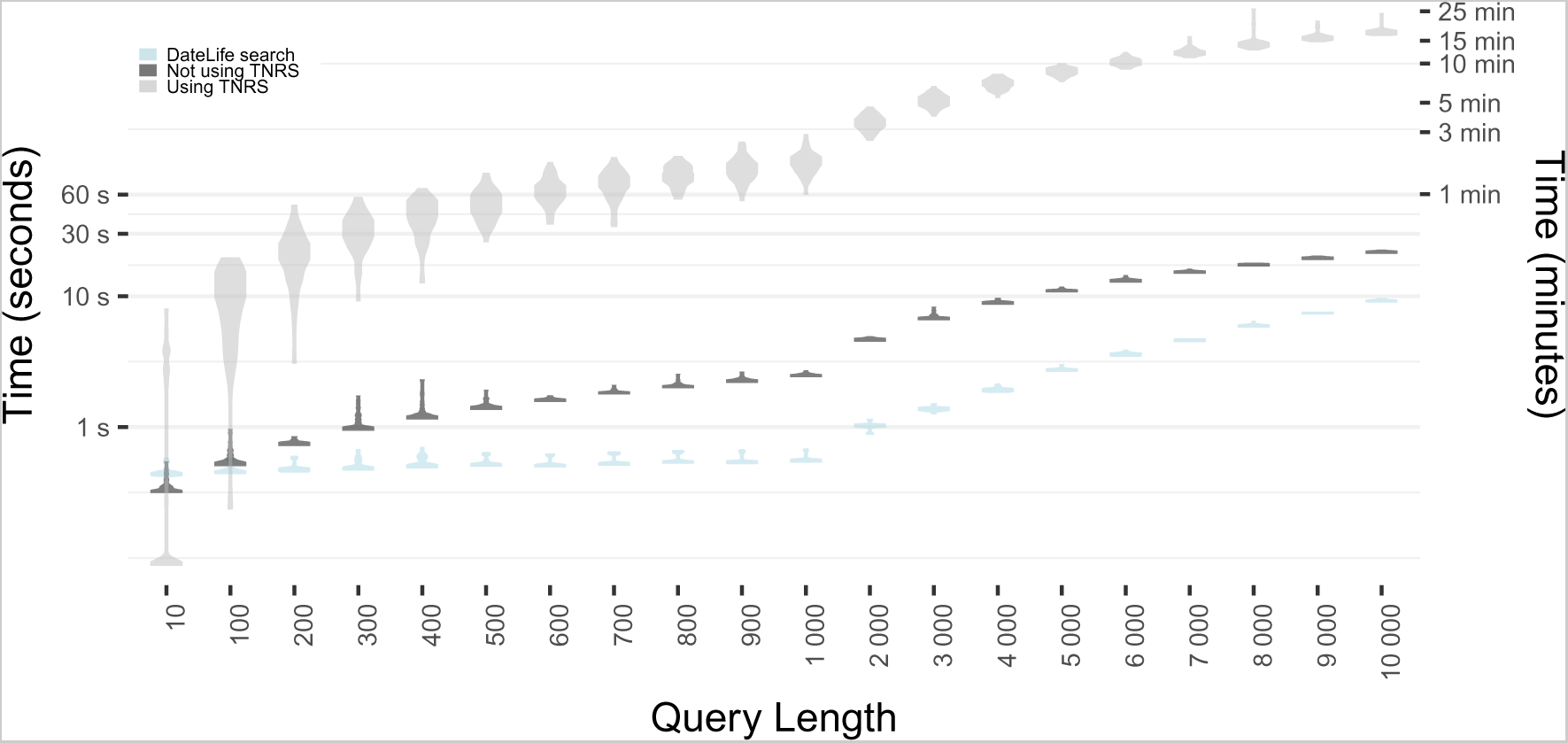
DateLife’s benchmarking results. Computation time used to process a query and a search across `datelife`’s chronogram database, relative to number of input taxon names. For each N = {10, 100, 200, …, 1 000, …, 9 000, 10 000}, we sampled N species names from the class Aves a hundred times, and then performed a datelife search processing the input names with Taxon Names Resolution Service (TNRS; light gray), and without processing names (dark gray). For comparison, we performed a search using an input that had been pre-processed with TNRS (light blue).

`datelife`’s code performance was evaluated with a set of unit tests designed and implemented with the R package testthat (R Core Team, 2018) that were run both locally with the devtools package (R Core Team, 2018), and on a public server using the continuous integration tool of GitHub actions (https://docs.github.com/en/actions). At present, unit tests cover more than 40% of `datelife`’s code (https://codecov.io/gh/phylotastic/datelife). Unit testing helps identify potential issues as code is updated or, more critically, as services code relies upon may change.

## Case studies

We illustrate the DateLife workflow using a family within the Passeriform birds encompassing the true finches, Fringillidae, as case study. The first example analyses 6 bird species and shows all steps of the workflow The second example is an analysis of 289 species in the family Fringillidae that are included in the NCBI taxonomy.

### A small example

#### Creating a search query

We chose 6 bird species within the Passeriformes. The sample includes two species of cardinals: the black-thighed grosbeak – *Pheucticus tibialis* and the crimson-collared grosbeak – *Rhodothraupis celaeno*; three species of buntings: the yellowhammer – *Emberiza citrinella*, the pine bunting – *Emberiza leucocephalos* and the yellow-throated bunting – *Emberiza elegans*; and one species of tanager, the vegetarian finch – *Platyspiza crassirostris*. Processing of input names found that *Emberiza elegans* is synonym for *Schoeniclus elegans* in the default reference taxonomy (OTT v3.3, June 1, 2021). For a detailed discussion on the state of the synonym, refer to Avibase (Avibase, 2022; Lepage, 2004; Lepage, Vaidya, & Guralnick, 2014). Discovering this synonym allowed assigning five age data points for the parent node of *Emberiza elegans*, shown as *Schoeniclus elegans* in figure 3A, which would not have had any data otherwise.

**Figure 3.**
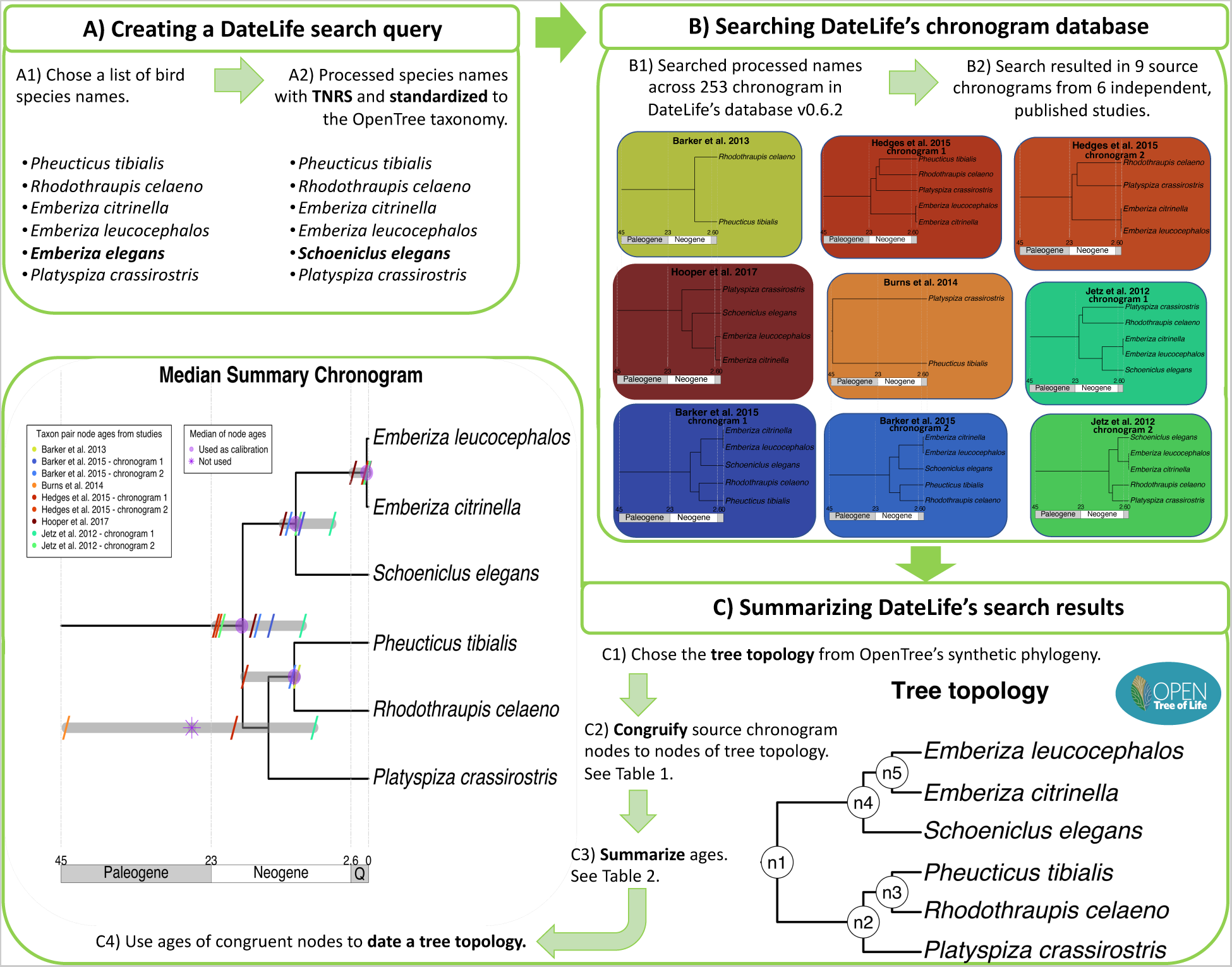
DateLife analysis results for a small sample of A) 6 bird species within the Passeriformes. B) Processed species names were found across 9 chronograms within 6 independent studies (Barker et al. (2012), Barker et al. (2015), Burns et al. (2014), Hedges et al. (2015), Hooper and Price (2017), Jetz et al. (2012).) C) This revealed 28 source age data points for the queried species names. Summarized age data is used as secondary calibrations to date a tree topology obtained from OpenTree’s synthetic tree, resulting in a summary chronogram of source ages.

#### Searching the database

DateLife used the processed input names to search the local chronogram database and found 9 matching chronograms in 6 different studies (Fig. 3B). Three studies matched five input names (Barker, Burns, Klicka, Lanyon, & Lovette, 2015; Hedges, Marin, Suleski, Paymer, & Kumar, 2015; Jetz, Thomas, Joy, Hartmann, & Mooers, 2012), one study matched four input names (Hooper & Price, 2017) and two studies matched two input names (Barker, Burns, Klicka, Lanyon, & Lovette, 2013; Burns et al., 2014). No studies matched all input names. Together, source chronograms provide 28 unique age data points, covering all nodes on our chosen tree topology to date (Table 1).

**Table 1.** Ages of congruified nodes. See Figure 3, step C2.

|  | Node Name | taxon A | taxon B | Node Age | Study chronogram |
| --- | --- | --- | --- | --- | --- |
| 1 | n1 | *Emberiza citrinella* | *Platypiza crassirostris* | 9.6509881 | Jetz et al. 2012 – chronogram 1 |
| 2 | n1 | *Pheucticus tibialis* | *Schoeniclus elegans* | 14.3336520 | Barker et al. 2015 – chronogram 1 |
| 3 | n1 | *Rhodothraupis celaeno* | *Schoeniclus elegans* | 16.2984859 | Barker et al. 2015 – chronogram 2 |
| 4 | n1 | *Emberiza citrinella* | *Platypiza crassirostris* | 16.9499615 | Hooper et al. 2017 |
| 5 | n1 | *Rhodothraupis celaeno* | *Schoeniclus elegans* | 21.5140867 | Jetz et al. 2012 – chronogram 2 |
| 6 | n1 | *Emberiza leucocephalos* | *Platypiza crassirostris* | 22.0000000 | Hedges et al. 2015 – chronogram 2 |
| 7 | n1 | *Emberiza citrinella* | *Platypiza crassirostris* | 22.3757277 | Hedges et al. 2015 – chronogram 1 |
| 8 | n2 | *Platypiza crassirostris* | *Rhodothraupis celaeno* | 7.9691925 | Jetz et al. 2012 – chronogram 1 |
| 9 | n2 | *Platypiza crassirostris* | *Rhodothraupis celaeno* | 19.7085830 | Jetz et al. 2012 – chronogram 2 |
| 10 | n2 | *Platypiza crassirostris* | *Rhodothraupis celaeno* | 19.7085900 | Hedges et al. 2015 – chronogram 2 |
| 11 | n2 | *Platypiza crassirostris* | *Rhodothraupis celaeno* | 19.7128363 | Hedges et al. 2015 – chronogram 1 |
| 12 | n2 | *Pheucticus tibialis* | *Platypiza crassirostris* | 44.2958603 | Burns et al. 2014 |
| 13 | n3 | *Pheucticus tibialis* | *Rhodothraupis celaeno* | 10.5304440 | Barker et al. 2015 – chronogram 1 |
| 14 | n3 | *Pheucticus tibialis* | *Rhodothraupis celaeno* | 10.5379092 | Barker et al. 2013 |
| 15 | n3 | *Pheucticus tibialis* | *Rhodothraupis celaeno* | 11.2095375 | Barker et al. 2015 – chronogram 2 |
| 16 | n3 | *Pheucticus tibialis* | *Rhodothraupis celaeno* | 18.1570685 | Hedges et al. 2015 – chronogram 1 |
| 17 | n4 | *Emberiza citrinella* | *Schoeniclus elegans* | 5.3238969 | Jetz et al. 2012 – chronogram 1 |
| 18 | n4 | *Emberiza leucocephalos* | *Schoeniclus elegans* | 9.8622460 | Barker et al. 2015 – chronogram 1 |
| 19 | n4 | *Emberiza leucocephalos* | *Schoeniclus elegans* | 10.3391445 | Jetz et al. 2012 – chronogram 2 |
| 20 | n4 | *Emberiza leucocephalos* | *Schoeniclus elegans* | 11.7317630 | Barker et al. 2015 – chronogram 2 |
| 21 | n4 | *Emberiza citrinella* | *Schoeniclus elegans* | 12.5133870 | Hooper et al. 2017 |
| 22 | n5 | *Emberiza citrinella* | *Emberiza leucocephalos* | 0.1407015 | Jetz et al. 2012 – chronogram 1 |
| 23 | n5 | *Emberiza citrinella* | *Emberiza leucocephalos* | 0.1516230 | Hedges et al. 2015 – chronogram 2 |
| 24 | n5 | *Emberiza citrinella* | *Emberiza leucocephalos* | 0.2011990 | Barker et al. 2015 – chronogram 1 |
| 25 | n5 | *Emberiza citrinella* | *Emberiza leucocephalos* | 0.2409300 | Barker et al. 2015 – chronogram 2 |
| 26 | n5 | *Emberiza citrinella* | *Emberiza leucocephalos* | 0.2732460 | Jetz et al. 2012 – chronogram 2 |
| 27 | n5 | *Emberiza citrinella* | *Emberiza leucocephalos* | 0.5760260 | Hedges et al. 2015 – chronogram 1 |
| 28 | n5 | *Emberiza citrinella* | *Emberiza leucocephalos* | 2.2898230 | Hooper et al. 2017 |

#### Summarizing search results

DateLife obtained OpenTree’s synthetic tree topology for these taxa (Fig. 3C), and congruified and mapped age data to nodes in this chosen topology (Table 1). The name processing step allowed including five data points for node “n4” (parent of *Schoeniclus elegans*; Fig. 3A) that would not have had any data otherwise due to name mismatch. Age summary statistics per node were calculated (Table 2) and used as calibrations to date the tree topology using the BLADJ algorithm. As expected, more inclusive nodes (e.g., node “n1”) have more variance in age data than less inclusive nodes (e.g., node “n5”). Summary age data for node “n2” were excluded as final calibration because they are older than age data of the more inclusive node, “n1” (Fig. 3C4).

**Table 2.** Summary of congruified nodes ages. See Figure 3, step C3.

| Node Name | Min Age | Q1 | Median Age | Mean Age | Q3 | Max Age | Variance | SD |
| --- | --- | --- | --- | --- | --- | --- | --- | --- |
| n1 | 9.6509881 | 15.316069 | 16.94996 | 17.5889860 | 21.757043 | 22.375728 | 22.2431847 | 4.7162681 |
| n2 | 7.9691925 | 19.708583 | 19.70859 | 22.2790124 | 19.712836 | 44.295860 | 177.3279940 | 13.3164558 |
| n3 | 10.5304440 | 10.536043 | 10.87372 | 12.6087398 | 12.946420 | 18.157069 | 13.7831237 | 3.7125630 |
| n4 | 5.3238969 | 9.862246 | 10.33914 | 9.9540875 | 11.731763 | 12.513387 | 7.8263782 | 2.7975665 |
| n5 | 0.1407015 | 0.176411 | 0.24093 | 0.5533641 | 0.424636 | 2.289823 | 0.6079318 | 0.7796998 |

### An example with the family of true finches

#### Creating a query

To obtain ages for all species within the family of true finches, Fringillidae, we ran a DateLife query using the “get species from taxon” flag, which gets all recognized species names within a named group from a taxonomy of choice. Following the NCBI taxonomy, our DateLife query has 289 Fringillidae species. This taxon-constrained approach implies that the final results of a full DateLife analysis will be done using a tree topology and ages for the species in a named group, which do not necessarily correspond to a monophyletic group. Users can change this behaviour by providing a monophyletic tree as input for a DateLife search, or as a tree topology for a DateLife summary.

#### Searching the database

Next, we used the processed species names in our DateLife query to identify chronograms with at least two Fringillidae species. The DateLife search identified 13 chronograms containing at least two Fringillidae species, published in 9 different studies (Barker et al., 2013, 2015; Burns et al., 2014; Claramunt & Cracraft, 2015; Gibb et al., 2015; Hedges et al., 2015; Hooper & Price, 2017; Jetz et al., 2012; Price et al., 2014). Once identified, DateLife pruned matching chronograms to keep Fringillidae species names on tips only, and transformed these pruned chronograms to pairwise distance matrices, revealing 1, 206 different age data points available for species within the Fringillidae (Supplementray Table S1).

#### Summarizing search results

The final step is to congruify and summarize the age data available for the Fringillidae species into single summary chronograms, using different types of summary ages, median and SDM. As explained in the “Description” section, a tree topology to summarize age data upon is required. By default, DateLife uses the topology from OpenTree’s synthetic tree that contains the species in the search query to summarize age data upon. According to OpenTree’s synthetic tree, species belonging to the family Fringillidae do not form a monophyletic group (Fig. 4).

**Figure 4.**
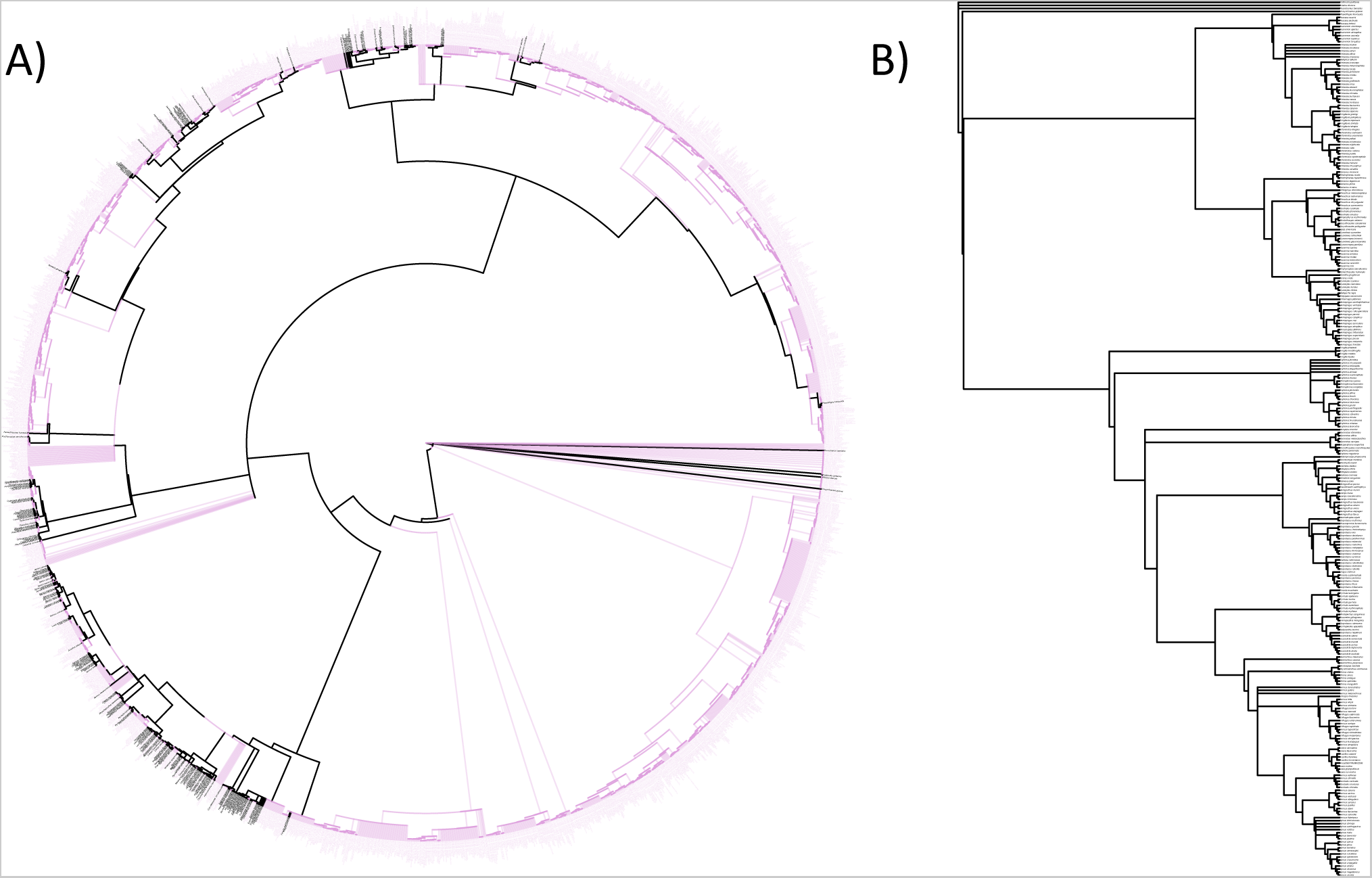
Tree topologies extracted from Open Tree of Life’s (OpenTree) synthetic phylogenetic tree. A) Topology of 2,333 tips and 1,305 internal nodes, encompassing bird species within the family Fringillidae following the NCBI taxonomy (black), as well as all other bird species that share the same Most Recent Common Ancestor (MRCA) node in OpenTree’s synthetic tree (purple). B) Subtree topology of of 289 tips and 253 internal nodes, resulting from pruning species that do not belong to the family Fringillidae according to the NCBI taxonomy (purple branches in topology A). Bird species within the Fringillidae are paraphyletic (Alström et al. 2014, Barker, Cibois, Schikler, Feinstein, & Cracraft 2004, Barker et al. 2013, Barker 2014, Barker et al. 2015, Beresford, Barker, Ryan, & Crowe 2005, Bryson Jr et al. 2014, Burleigh, Kimball, & Braun 2015, Burns et al. 2014, Chaves, Hidalgo, & Klicka 2013, Claramunt & Cracraft 2015, Gibb et al. 2015, Hackett et al. 2008, Jetz et al. 2012, Johansson, Fjeldså, & Bowi 2008, Kimball et al. 2019, Klicka et al. 2014, Lamichhaney et al. 2015, Lerner, Meyer, James, Hofreiter, & Fleischer 2011, Lovette et al. 2010, Moyle et al. 2016, Ödeen, Håstad, & Alström 2011, Oliveros et al. 2019, Päckert et al. 2012, Parchman, Benkman, & Mezquida 2007, Powell et al. 2014, Price et al. 2014, Pulgarín-R, Smith, Bryson Jr, Spellman, & Klicka 2013, Selvatti, Gonzaga, & Moraes Russo 2015, Tietze, Päckert, Martens, Lehmann, & Sun 2013, Treplin et al. 2008, Zuccon, Prŷs-Jones, Rasmussen, & Ericson 2012).

Age data from source chronograms was congruified to OpenTree’s topology (Fig. 4B), reducing the age data set to 818 different data points (Supplementray Table S2). For each congruent node, age summary statistics were calculated and used as fixed secondary calibrations over the chosen tree topology, to obtain a fully dated phylogeny with the program BLADJ (Fig. 5).

**Figure 5.**
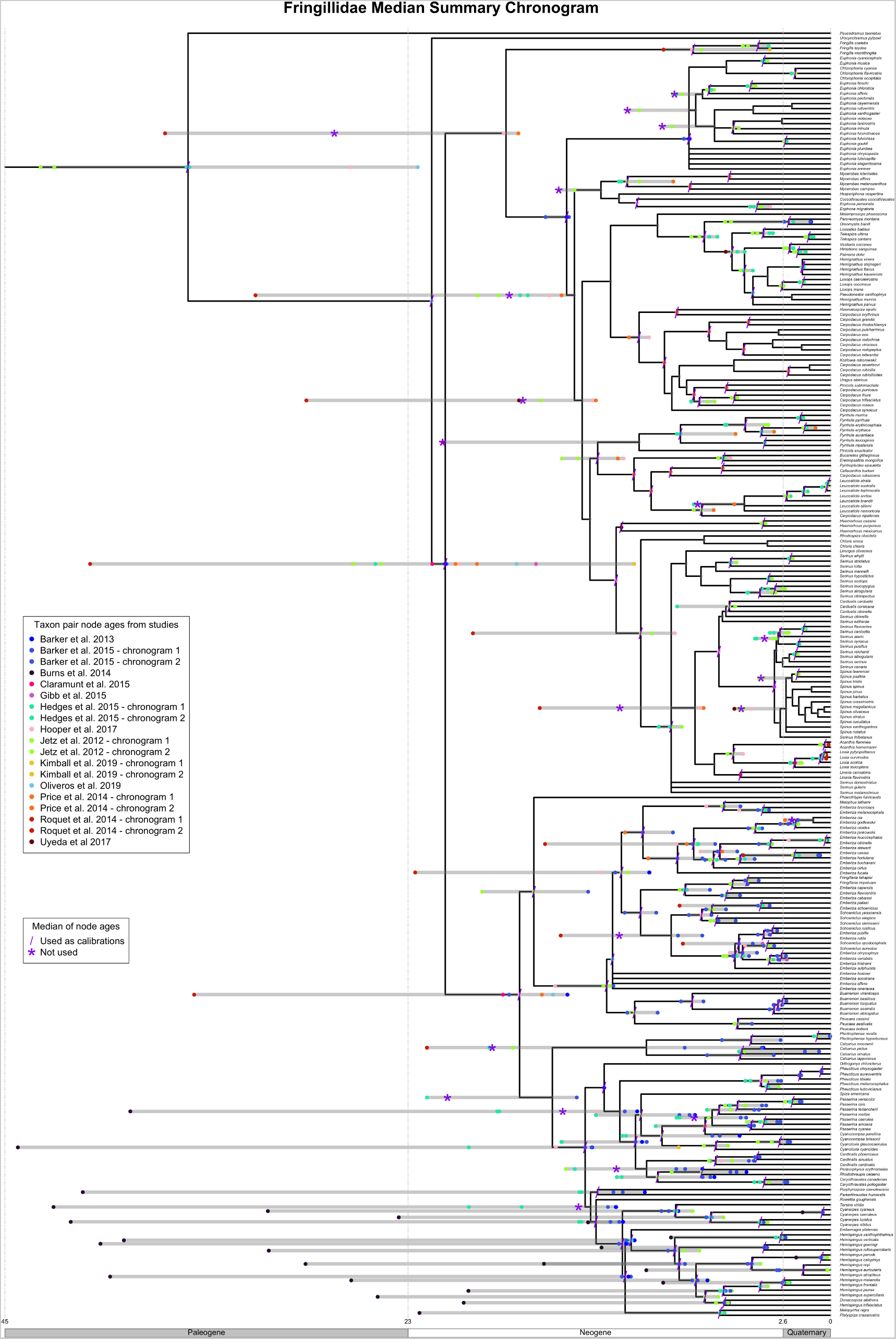
Fringillidae median summary chronogram generated with DateLife. It has 256 tips and 233 nodes, from which 212 have age data from at least one published chronogram.

## Cross-validation test

We performed a cross validation analysis of the DateLife workflow using the Fringillidae chronograms. We used the individual tree topologies from each of the 19 source chronograms from 13 studies as inputs, treating their node ages as unknown. We then estimated dates for these topologies using the node ages from the chronograms from the other studies as calibrations and smoothing using BLADJ. We found that node ages from original study, and ages estimated using all other age data available are correlated (Fig. 6). For five studies, Datelife tended to underestimate ages for topologically deeper nodes (those with many descendant taxa, aka ‘closer to the root’) relative to the original estimate, and overestimate ages for nodes closer to the tips. Accordingly, root ages are generally older in the original study than estimated using cross-validated ages (Supplementary Fig. S1).

**Figure 6.**
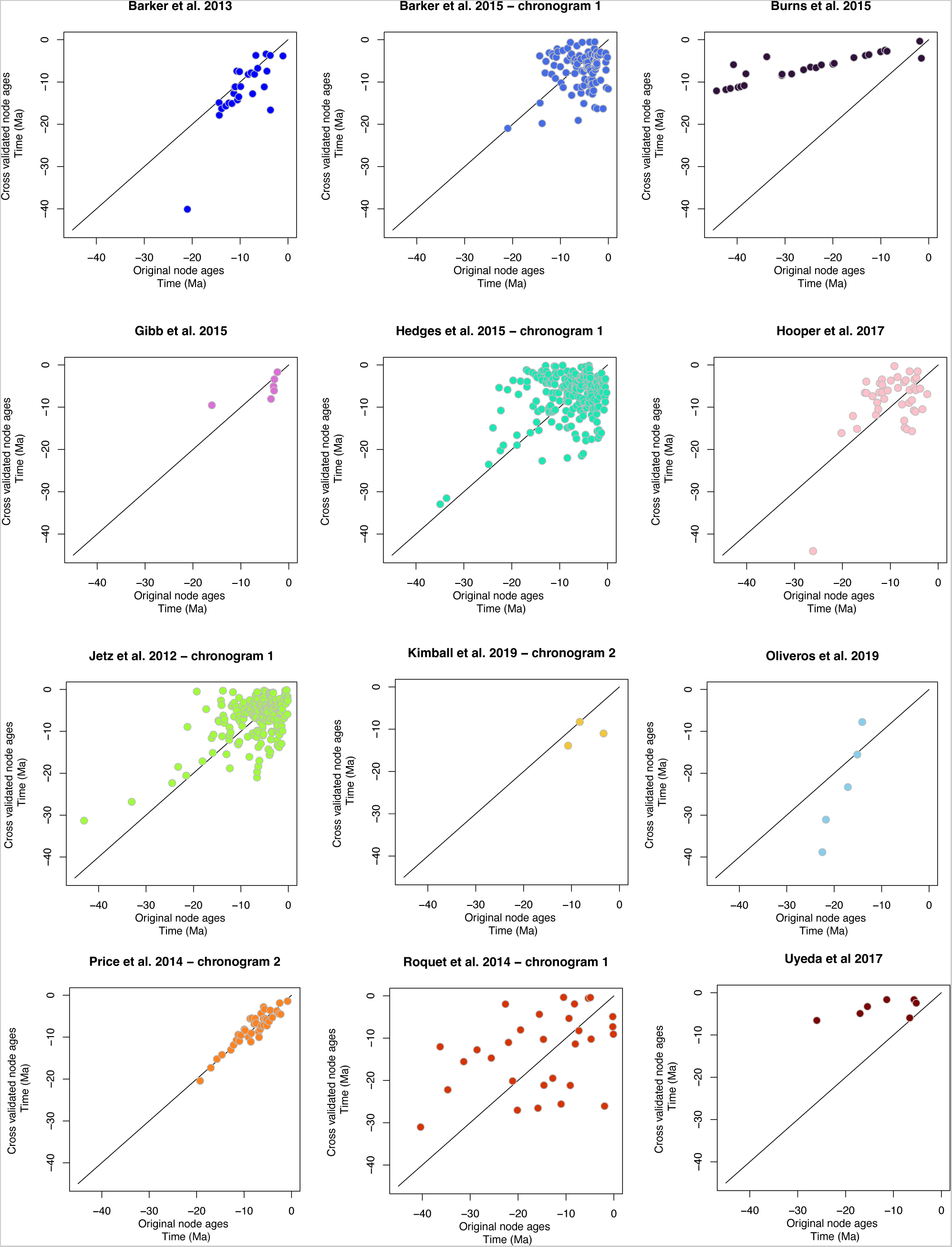
Results from cross validation analysis. Each plot compares the original age estimate (x axis) with the age obtained with a DateLife analysis (y axis), per node.

## Discussion

DateLife makes state-of-the-art data on evolutionary time frame easily accessible for comparison, reuse, and reanalysis, to researchers in all areas of science and with all levels of expertise in the matter. It is an open service that does not require any expert biological knowledge from users –besides the names of the species or group they want to work with, for any of its functionality.

A total of 99,474 unique terminal taxa are represented in DateLife’s database. Incorporation of more chronograms into the database will continue to improve DateLife’s services. One option to increase the number of chronograms in the DateLife database is the Dryad data repository. Methods to automatically mine chronograms from Dryad could be designed and implemented. However, Dryad’s metadata system has no information to automatically detect branch length units, and those would still need to be determined manually by a human curator. We would like to emphasize on the importance of sharing chronogram data, including systematically curated metadata, into open repositories, such as OpenTree’s Phylesystem (McTavish et al., 2015) for the benefit of the scientific community as a whole.

### Age variation in source chronograms

Conflict in estimated ages among alternative studies is common in the literature. See, for example, the robust ongoing debate about crown group age of angiosperms (Barba-Montoya, Reis, Schneider, Donoghue, & Yang, 2018; Magallón, Gómez-Acevedo, Sánchez-Reyes, & Hernández-Hernández, 2015; Ramshaw et al., 1972; Sanderson & Doyle, 2001; Sauquet, Ramírez-Barahona, & Magallón, 2021). Source chronograms available for the same organisms have potentially been estimated implementing calibrations very differently. For example, the chronograms from Burns et al. (2014) were inferred using molecular substitution rate estimates across birds (Weir & Schluter, 2008), and have much older age estimates for the same nodes than chronograms that were inferred using fossils as calibrations (Figs. 5, 6; Supplementary Figs. S1, S5).

Different calibration implementations might also imply fundamentally distinct evolutionary hypotheses (Antonelli et al., 2017). For example, two independent researchers working on the same clade should both carefully select and justify their choices of fossil calibration placement. Yet, if one researcher concludes that a fossil should calibrate the ingroup of a clade, while another researcher concludes that the same fossil should calibrate the outgroup of the clade, the resulting age estimates will differ, as the placement of calibrations as stem or crown group has been proven to significantly affect time of lineage divergence estimates (Sauquet, 2013).

#### Primary vs Secondary calibrations

While most chronograms in DateLife’s database are constructed using primary calibrations (molecular substitution rates or ages obtained from the fossil record or geological events), DateLife summarizes chronograms using secondary calibrations (ages coming from other chronograms). Graur and Martin (2004) cautioned on the increased error and uncertainty in estimated ages when using secondary calibrations in dating analyses. Schenk (2016) showed that, in simulations, divergence times inferred using secondary calibrations are significantly younger than those inferred with primary calibrations, when obtained with Bayesian inference methods, and when priors are implemented in similar ways in both analyses. Accordingly, the scientific community seems to have more confidence in chronograms obtained from a single analysis, using fossil data as primary sources of calibrations (Schenk, 2016), and using fossils that have been widely discussed and curated as calibrations to date other trees, making sure that all data reflect a coherent evolutionary history (Sauquet, 2013), as for example done by Antonelli et al. (2017). There have been attempts to create fossil calibration databases (Ksepka et al., 2015), though these still have room to grow.

It seems that using several (as opposed to just a few) secondary calibrations can provide sufficient information to alleviate or even neutralize potential biases (Sauquet, 2013). Certainly, further studies are required to fully understand the effect of secondary calibrations on outputs from different tree dating methods, and on downstream analyses. It is possible that secondary calibrations can be safely used with dating methods that do not require setting priors, such as penalized likelihood (Sanderson, 2003), with methods that do not make any assumptions on the ages and fix them to a node on a tree topology, such as BLADJ (Webb et al., 2008; Webb & Donoghue, 2005), or methods that summarize age data unto a tree topology.

Our cross validation analysis might provide some insight in this regard. When ages are estimated with secondary calibrations, nodes closer to the root do tend to be slightly younger than ages estimated with primary calibrations. However, nodes closer to the tip tend to be older when estimated using secondary calibrations with a dating method that does not make any prior assumptions on the nature of the calibrations themselves (Supplementary Figures S2-S20). The only excpetion to tjis was observed on results of the cross validation analysis of the Burns et al. (2014) chronogram, which results in much younger node ages when estimated using secondary calibrations (Supplementary Figs. S1, S5).

### Sumarizing chronograms

By default, DateLife currently summarizes all source chronograms that overlap with at least two species names. Users can exclude source chronograms if they have reasons to do so. Strictly speaking, a good chronogram should reflect the real time of lineage divergence accurately and precisely. To our knowledge, there are no tested measures to determine independently when a chronogram is better than another. Yet, several characteristics of the data used for dating analyses, as well as from the output chronogram itself, could be used to score the quality of source chronograms.

Some measures that have been proposed are the proportion of lineage sampling and the number of calibrations used Magallón et al. (2015). Some characteristics that are often cited in published studies as a measure of improved age estimates as compared to previously published estimates are: quality of alignment (missing data, GC content), lineage sampling (strategy and proportion), phylogenetic and dating inference method, number of fossils used as calibrations, support for nodes and ages, and magnitude of confidence intervals. DateLife provides an opportunity to capture concordance and conflict among date estimates, which can also be used as a metric for chronogram reliability. Its open database of chronograms allows other researchers to do such analyses themselves reproducibly, and without needing permission. Though, of course, they should follow proper citation practices, especially for the source chronogram studies.

The exercise of summarizing age data from across multiple studies provides the opportunity to work with a more inclusive chronogram, that reflects a unified evolutionary history for a lineage, by putting together evidence from different hypotheses. The largest, and taxonomically broadest chronogram currently available from OpenTree was constructed summarizing age data from 2,274 published chronograms using NCBI’s taxonomic tree as backbone (Hedges et al., 2015). A summarizing exercise may also amplify the effect of uncertainty and errors in source data, and blur parts of the evolutionary history of a lineage that might only be reflected in source chronograms and lost on the summary chronogram (Sauquet et al., 2021).

### Effects on downstream analyses

For downstream analyses, using alternative chronogram may deeply affect our inferences (Title & Rabosky, 2016), particularly when studying phenomena dependent on the timing of species diversification events, such as macroevolutionary processes.

In ecology and conservation biology, incorporating at least some data on lineage divergence times represents a relevant improvement for testing alternative hypothesis using phylogenetic distance (Webb et al., 2008). Hence, DateLife’s workflow features different ways of estimating node ages in the absence of calibrations and branch length information for certain taxa. “Making up” branch lengths is a common practice in scientific publications: Jetz et al. (2012), created a chronogram of all 9, 993 bird species, where 67% had molecular data and the rest was simulated; Rabosky et al. (2018) created a chronogram of 31, 536 ray-finned fishes, of which only 37% had molecular data; Smith and Brown (2018) constructed a chronogram of 353, 185 seed plants where only 23% had molecular data.

Notably, risks come with this practice. Taken to the extreme, one could make a fully resolved, calibrated tree of all modern and extinct taxa using a single taxonomy and a single calibration, using polytomy resolution and branch estimation methods. There has yet to be a thorough analysis of what can go wrong when one extends inferences beyond the data in this way, so we urge caution; we also urge readers to follow the example of the large tree papers cited above, by carefully considering the statistical assumptions being made, and assessing the consistency of the results with prior work.

## Conclusions

Knowledge of the evolutionary time frame of organisms is key to many research areas: trait evolution, species diversification, biogeography, macroecology and more. It is also crucial for education, science communication and policy, but generating chronograms is difficult, especially for those who want to use phylogenies but who are not systematists, or do not have the time to acquire and develop the necessary knowledge and skills to construct them on their own. Importantly, years of primarily public funded research have resulted in vast amounts of chronograms that are already available on scientific publications, but hidden to the public and scientific community for reuse.

The DateLife project allows for easy and fast summary of public and state-of-the-art data on time of lineage divergence. It provides a straightforward way to get an informed idea on the state of knowledge of the time frame of evolution of different regions of the tree of life, and allows identification of regions that require more research, or that have conflicting information. It is available as an R package, and as a web-based R shiny application at www.datelife.org Both summary and newly generated trees are useful to evaluate evolutionary hypotheses in different areas of research. The DateLife project helps with awareness of the existing variation in expert time of divergence data, and will foster exploration of the effect of alternative divergence time hypothesis on the results of analyses, nurturing a culture of more cautious interpretation of evolutionary results.

## Availability

The DateLife software is free and open source and it can be used through its R shiny web application at http://www.datelife.org, through the `datelife` R package, and through Phylotastic’s project web portal https://phylo.cs.nmsu.edu/. DateLife’s web application is maintained using RStudio’s shiny server and the shiny package open infrastructure, as well as Docker and OpenTree’s infrastructure (dates.opentreeoflife.org/datelife). `datelife`’s R package stable version is available for installation from the CRAN repository (https://cran.r-project.org/package=datelife) using the command `install.packages(pkgs = “datelife”)` from within R. Development versions are available from the GitHub repository (https://github.com/phylotastic/datelife) and can be installed using the command `devtools::install_github(“phylotastic/datelife”)`.

## Supplementary Material

Code used to generate all versions of this manuscript, the biological examples, as well as the benchmark of functionalities are available at datelifeMS1, datelife_examples, and datelife_benchmark repositories in LLSR’s GitHub account.

## Funding

Funding was provided by the US National Science Foundation (NSF) grants ABI-1458603 to the Datelife project; DBI-0905606 to the National Evolutionary Synthesis Center (NESCent), ABI-1458572 to the Phylotastic project, and ABI-1759846 to the Open Tree of Life project.

## Acknowledgements

The DateLife project was born as a prototype tool aiming to provide these services, and was initially developed over a series of hackathons at the National Evolutionary Synthesis Center, NC, USA (Stoltzfus et al., 2013). We thank colleagues from the O’Meara Lab at the University of Tennesse Knoxville for suggestions, discussions and software testing. The late National Evolutionary Synthesis Center (NESCent), which sponsored hackathons that led to initial work on this project. The team that assembled DateLife’s first proof of concept: Tracy Heath, Jonathan Eastman, Peter Midford, Joseph Brown, Matt Pennell, Mike Alfaro, and Luke Harmon. The Open Tree of Life project that provides the open, metadata rich repository of trees used to construct DateLife’s chronogram database. The many scientists who publish their chronograms in an open, reusable form, and the scientists who curate them for deposition in the Open Tree of Life repository. The NSF for funding nearly all the above, in addition to the ABI grant that funded this project itself.

## Notes

http://datelife.opentreeoflife.org/query/

https://github.com/phylotastic/datelife

https://github.com/LunaSare/datelife_examples

https://github.com/LunaSare/datelifeMS1

### Competing Interest Statement

The authors have declared no competing interest.

### Summary of Updates

Adding a new co-author; updating and adding examples; updating all figures and tables; a more thorough methodology; synthesizing discussion.

