## Supplementary figures and images for "DateLife: leveraging databases and analytical tools to reveal the dated Tree of Life"

### Supplementary Figure S

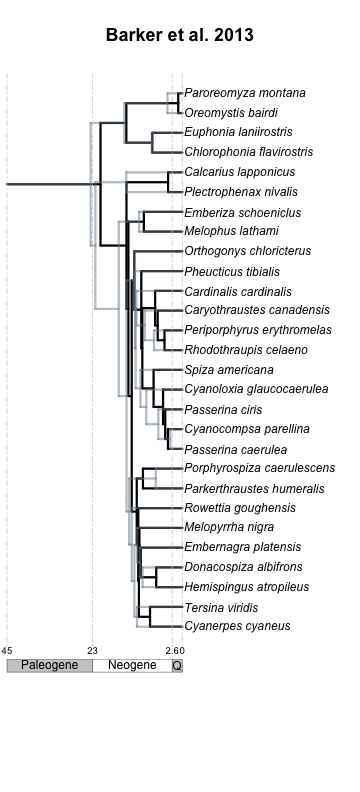

### Supplementary Figure S

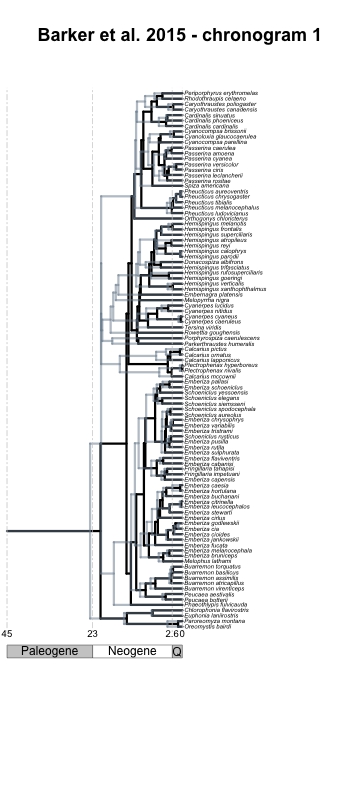

### Supplementary Figure S

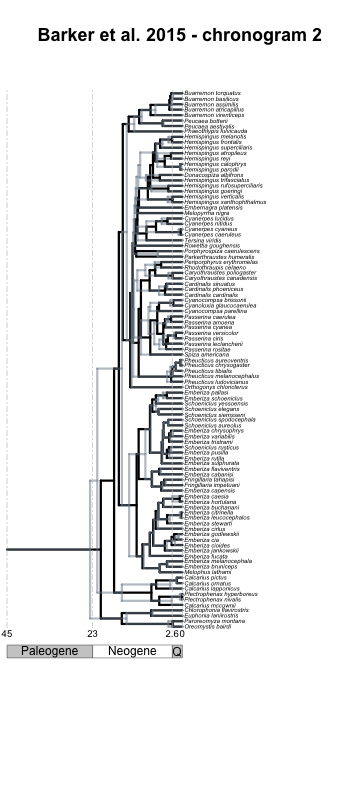

### Supplementary Figure S

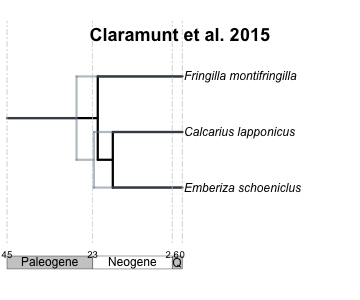

### Supplementary Figure S

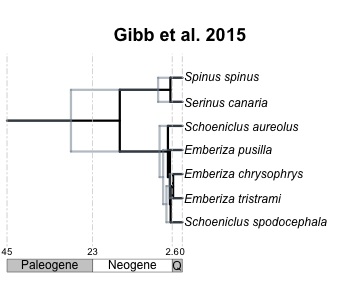

### Supplementary Figure S

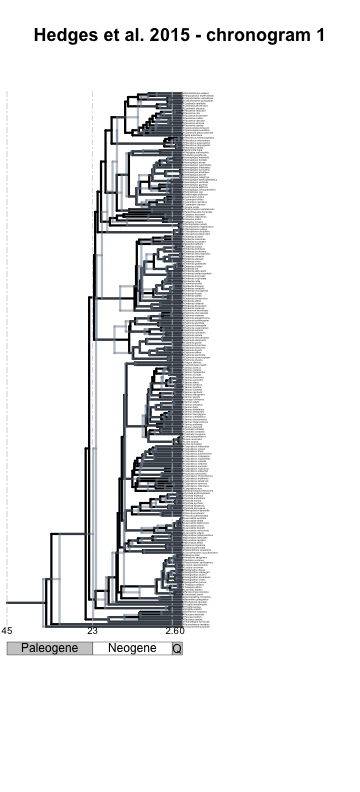

### Supplementary Figure S

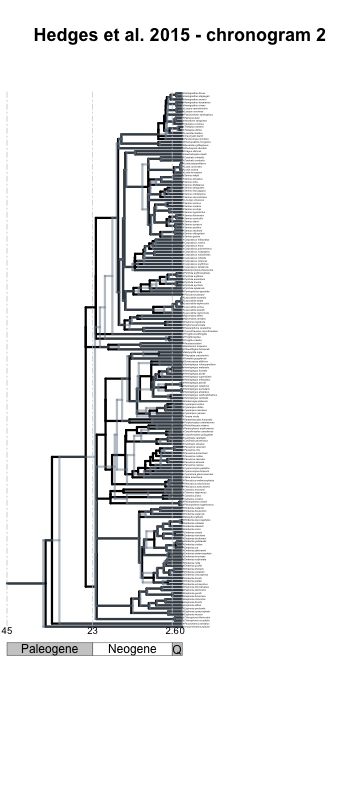

### Supplementary Figure S

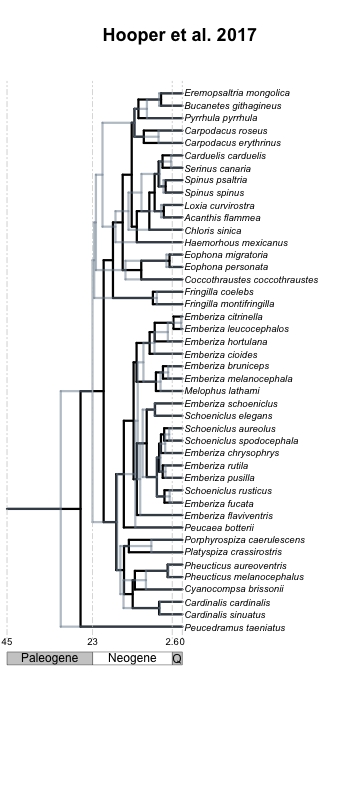

### Supplementary Figure S

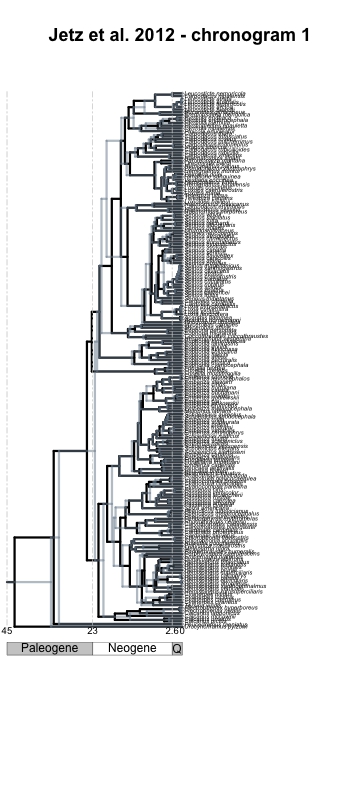

### Supplementary Figure S

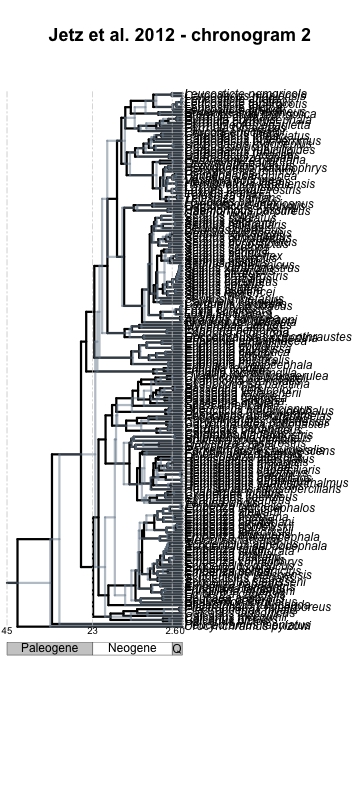

### Supplementary Figure S

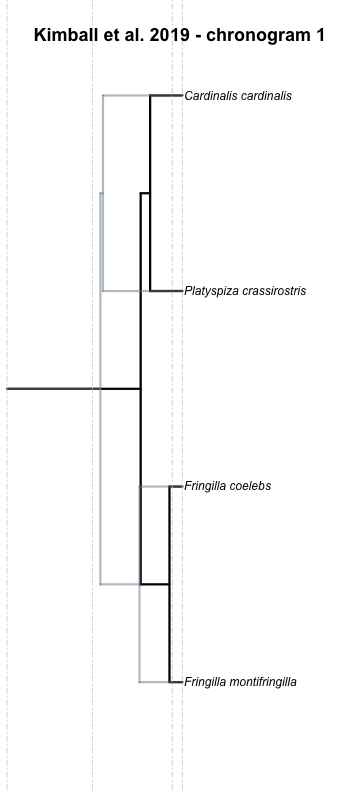

### Supplementary Figure S

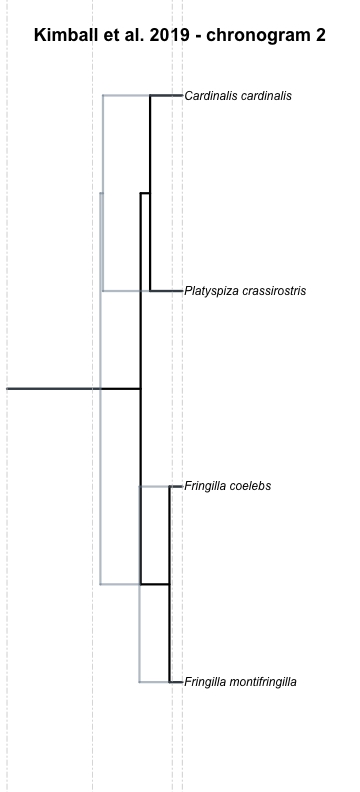

### Supplementary Figure S

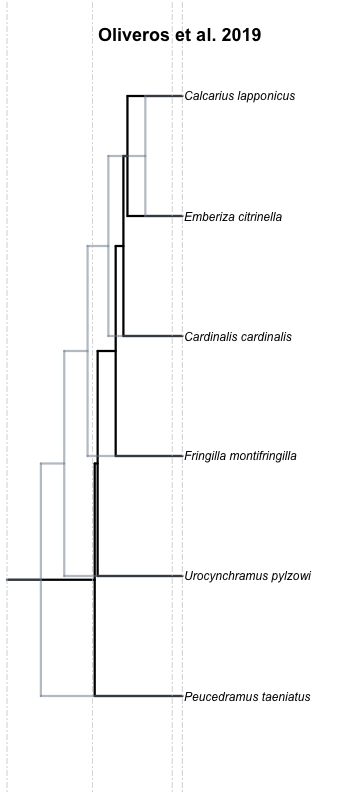

### Supplementary Figure S

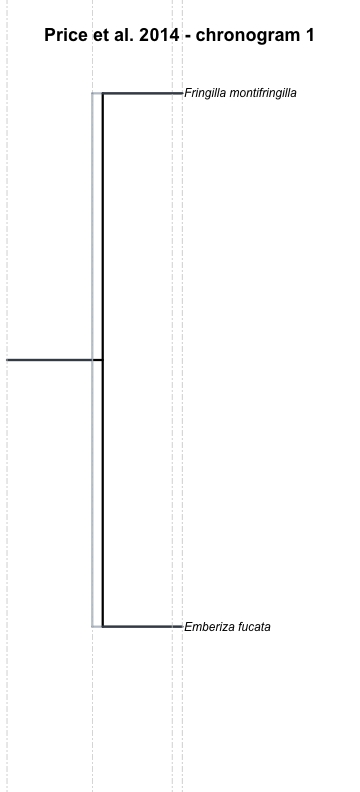

### Supplementary Figure S

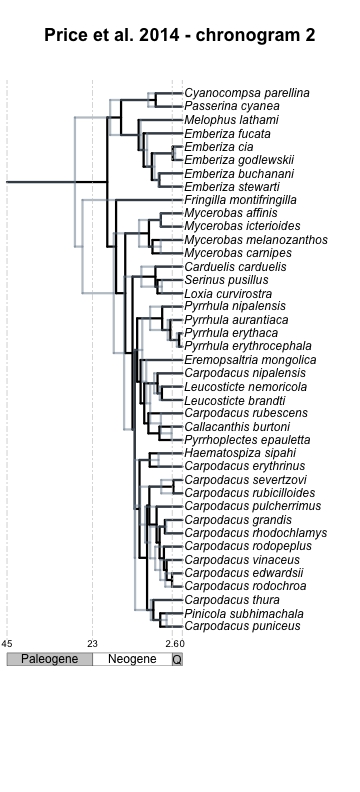

### Supplementary Figure S

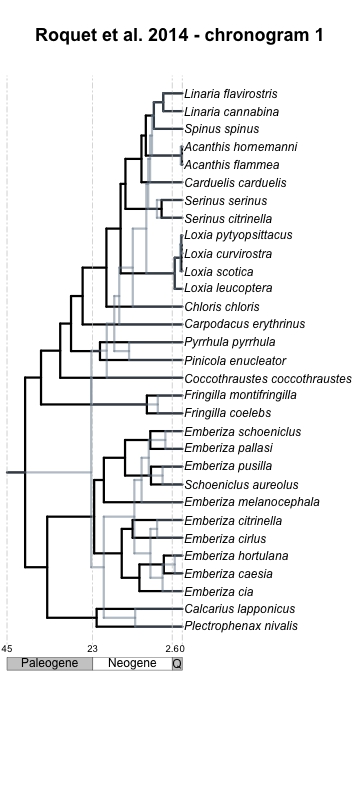

### Supplementary Figure S

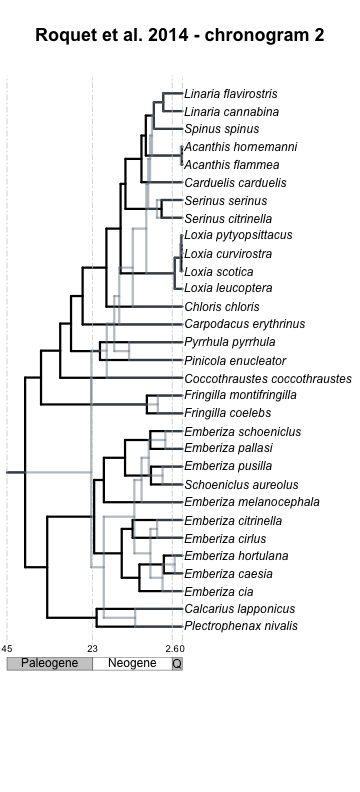

### Supplementary Figure S

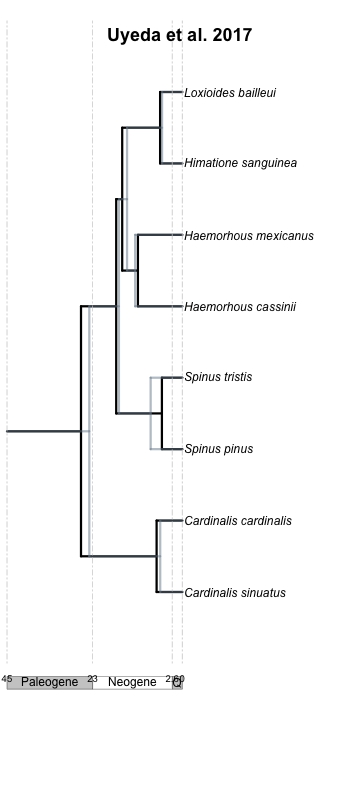

### Supplementary Figure S1

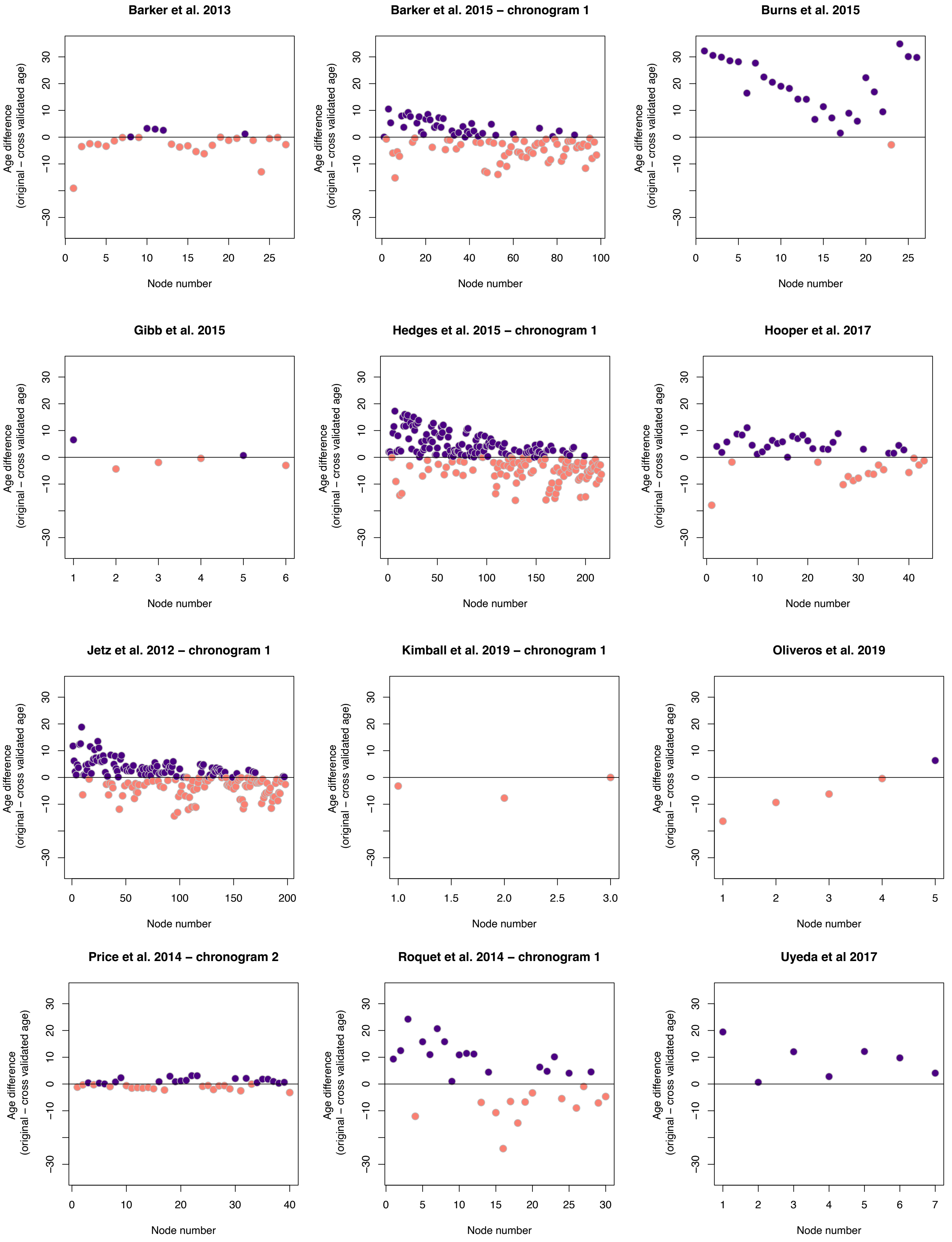
