## SupplementaryTable S2 for "DateLife: leveraging databases and analytical tools to reveal the dated Tree of Life"

| Node Name | Min Age | Q1 | Median Age | Mean Age | Q3 | Max Age | Variance | SD |
|---|---|---|---|---|---|---|---|---|
| n257 | 22.4922461 | 28.3308351 | 35.0195851 | 34.0041747 | 40.5051139 | 43.0587563 | 6.951043e+01 | 8.337291261 |
| n258 | 21.7398085 | 21.7398085 | 21.7398085 | 21.7398085 | 21.7398085 | 21.7398085 | NA | NA |
| n259 | 10.6957998 | 19.4987850 | 20.9992411 | 22.5521354 | 24.8109359 | 40.3552933 | 6.196330e+01 | 7.871676844 |
| n260 | 14.3336515 | 15.1266340 | 16.9499603 | 20.1348914 | 17.8493180 | 34.6833663 | 6.965831e+01 | 8.346155338 |
| n261 | 13.8254294 | 17.3500521 | 20.8746749 | 18.9000348 | 21.4373374 | 22.0000000 | 1.963030e+01 | 4.430609873 |
| n262 | 8.2674570 | 12.9942327 | 13.3766918 | 17.1809206 | 19.7085900 | 44.2958603 | 1.201132e+02 | 10.959617046 |
| n263 | 11.6676580 | 11.9166862 | 13.7253610 | 18.3018211 | 18.2685746 | 42.3492073 | 1.215023e+02 | 11.022806779 |
| n264 | 11.3074645 | 11.6411873 | 12.7840747 | 17.2544161 | 13.6774289 | 41.4154443 | 1.411443e+02 | 11.880415266 |
| n265 | 11.0467304 | 11.1243986 | 11.3445148 | 18.3814487 | 18.6015648 | 39.7900348 | 2.037460e+02 | 14.273962041 |
| n266 | 10.8823208 | 10.9826962 | 11.1967141 | 18.1343797 | 18.3483977 | 39.2617699 | 1.984289e+02 | 14.086477980 |
| n267 | 5.7738060 | 5.9827515 | 6.0261000 | 9.2429048 | 6.0261040 | 22.4057625 | 5.415493e+01 | 7.359003149 |
| n268 | 10.6775303 | 10.7242787 | 10.8657943 | 15.4566849 | 11.0696627 | 38.5162645 | 1.276461e+02 | 11.298059475 |
| n269 | 7.1433202 | 7.9864380 | 8.4893522 | 11.3687348 | 8.6779130 | 30.6197691 | 7.236208e+01 | 8.506590153 |
| n270 | 7.9299083 | 8.0911383 | 8.2523682 | 14.9318196 | 18.4327753 | 28.6131824 | 1.404108e+02 | 11.849504652 |
| n271 | 6.6427435 | 6.8390883 | 7.3322468 | 10.2785734 | 7.5080840 | 26.1228047 | 6.038528e+01 | 7.770796804 |
| n272 | 6.1123530 | 6.4730706 | 6.8337881 | 12.5423463 | 15.7573429 | 24.6808977 | 1.106384e+02 | 10.518480889 |
| n273 | 5.5318420 | 5.5833590 | 5.7899205 | 8.6281640 | 6.2423575 | 19.9933411 | 4.044325e+01 | 6.359500683 |
| n274 | 5.3809230 | 5.5853838 | 5.8298120 | 7.7352529 | 6.0160288 | 19.7332102 | 2.805236e+01 | 5.296447891 |
| n275 | 3.4184700 | 3.7210793 | 3.8219495 | 3.7290430 | 3.8299133 | 3.8538030 | 4.309463e−02 | 0.207592454 |
| n276 | 2.2243040 | 2.4150750 | 2.5820770 | 3.4248889 | 2.6077595 | 9.1221722 | 6.334873e+00 | 2.516917323 |
| n277 | 3.8092620 | 4.2588651 | 4.2842073 | 6.1297245 | 4.4683045 | 15.6205824 | 2.167229e+01 | 4.655351214 |
| n278 | 0.1526050 | 0.1706170 | 1.4824100 | 2.7078835 | 2.8861148 | 10.0425071 | 1.465053e+01 | 3.827601636 |
| n279 | 0.1680660 | 0.1879020 | 0.4853190 | 0.5679572 | 0.5357770 | 1.8749574 | 3.610337e−01 | 0.600860835 |
| n280 | 3.4483380 | 3.5305280 | 3.6127180 | 6.5136472 | 8.0463018 | 12.4798856 | 2.670376e+01 | 5.167567689 |
| n281 | 2.4430780 | 2.6000780 | 2.7193950 | 3.5724223 | 2.9752185 | 8.6938904 | 5.140696e+00 | 2.267310259 |
| n282 | 7.5082385 | 8.3697974 | 8.4401100 | 11.2226005 | 8.9164799 | 30.6687844 | 6.197376e+01 | 7.872341292 |
| n283 | 5.8317240 | 6.4987639 | 6.5200370 | 8.8856028 | 6.6583505 | 23.5332301 | 4.180233e+01 | 6.465472280 |
| n284 | 3.4274700 | 3.7384500 | 3.8320100 | 5.0932213 | 3.8501610 | 13.2158472 | 1.285324e+01 | 3.585141636 |
| n285 | 0.4196550 | 0.4296950 | 0.4397350 | 0.7955567 | 0.9835076 | 1.5272802 | 4.016652e−01 | 0.633770635 |
| n286 | 10.1178768 | 11.4491148 | 12.7012298 | 16.8790191 | 13.6535585 | 40.7510339 | 1.386228e+02 | 11.773819838 |
| n287 | 12.3367949 | 12.3376283 | 12.3384617 | 12.3384617 | 12.3392951 | 12.3401285 | 5.556454e−06 | 0.002357213 |
| n288 | 10.5304438 | 11.2095374 | 14.5872601 | 16.8895312 | 18.0378600 | 38.1656654 | 7.405778e+01 | 8.605682847 |
| n289 | 10.0136627 | 10.2731275 | 10.6297969 | 12.0463515 | 14.6389300 | 14.6762402 | 5.730138e+00 | 2.393770654 |
| n290 | 6.9794359 | 8.1625350 | 11.6644360 | 11.1075354 | 14.1800830 | 14.4236400 | 1.080151e+01 | 3.286565213 |
| n291 | 6.3225445 | 6.7594660 | 9.4206630 | 9.1343196 | 11.5144940 | 11.6491100 | 6.174613e+00 | 2.484876938 |
| n292 | 2.3548040 | 2.7311138 | 3.6064205 | 3.3985392 | 4.0769365 | 4.1504700 | 6.397417e−01 | 0.799838568 |
| n293 | 4.5800958 | 5.2886520 | 7.0053400 | 6.8776178 | 8.6590715 | 8.6624420 | 3.321884e+00 | 1.822603506 |
| n294 | 4.1787990 | 5.1289133 | 6.2581125 | 5.7937331 | 6.6097345 | 6.6220000 | 1.034429e+00 | 1.017068728 |
| n295 | 3.8634520 | 3.9275823 | 3.9917125 | 3.9917125 | 4.0558428 | 4.1199730 | 3.290151e−02 | 0.181387739 |
| n296 | 7.3844932 | 7.9588171 | 10.3269032 | 10.2795400 | 12.7697076 | 12.7883343 | 6.333546e+00 | 2.516653743 |
| n297 | 4.9521150 | 6.2403953 | 9.1851225 | 8.6397538 | 11.2511962 | 11.3578560 | 7.757092e+00 | 2.785155592 |
| n298 | 5.5851610 | 5.9154510 | 6.2457410 | 6.2457410 | 6.5760310 | 6.9063210 | 8.727319e−01 | 0.934201195 |
| n299 | 2.1957830 | 2.2987040 | 2.7709670 | 2.9264263 | 3.3986893 | 3.9679880 | 6.835455e−01 | 0.826768107 |
| n300 | 4.3913853 | 5.6151566 | 6.8389280 | 6.9201044 | 8.1844640 | 9.5300000 | 6.606283e+00 | 2.570268969 |
| n301 | 4.5936013 | 5.3303791 | 7.4300749 | 6.8253582 | 8.2159191 | 8.3900503 | 3.013068e+00 | 1.735819148 |
| n302 | 4.1722035 | 4.2438610 | 4.3155185 | 4.3155185 | 4.3871760 | 4.4588335 | 4.107838e−02 | 0.202678017 |
| n303 | 2.7293880 | 3.0636078 | 3.6111525 | 3.5409692 | 3.7936123 | 4.5585970 | 4.402972e−01 | 0.663548926 |
| n304 | 3.9059560 | 4.4762974 | 6.0724853 | 5.6097295 | 6.7147436 | 6.7343860 | 1.749722e+00 | 1.322770739 |
| n305 | 3.4997120 | 3.9591913 | 5.0692245 | 4.8096333 | 5.6816643 | 5.7446480 | 1.024547e+00 | 1.012199088 |
| n306 | 1.5799340 | 1.7582140 | 2.0683305 | 2.0120555 | 2.1878660 | 2.4731850 | 1.128194e−01 | 0.335886015 |
| n307 | 2.6891060 | 3.1978420 | 3.7704640 | 3.7481452 | 4.3678785 | 4.6460055 | 5.876274e−01 | 0.766568573 |
| n308 | 2.8294780 | 3.2528065 | 3.4463515 | 3.3052428 | 3.4987878 | 3.4987900 | 1.030450e−01 | 0.321006208 |
| n309 | 1.6036030 | 1.6306588 | 1.6577145 | 1.6577145 | 1.6847703 | 1.7118260 | 5.856109e−03 | 0.076525217 |
| n310 | 0.4871060 | 0.4951133 | 0.5031205 | 0.5031205 | 0.5111278 | 0.5191350 | 5.129284e−04 | 0.022647923 |
| n311 | 3.6532999 | 7.9414692 | 18.4347739 | 14.6797389 | 18.5969520 | 21.9999770 | 5.554413e+01 | 7.452793385 |
| n312 | 3.0969730 | 4.0047720 | 4.9125710 | 4.9125710 | 5.8203700 | 6.7281690 | 6.592792e+00 | 2.567643315 |
| n313 | 0.6191110 | 2.2175475 | 4.8227280 | 3.6045113 | 4.9784123 | 4.9830400 | 4.119455e+00 | 2.029644004 |
| n314 | 2.7181595 | 3.5206588 | 4.3231580 | 4.3231580 | 5.1256573 | 5.9281565 | 5.152040e+00 | 2.269810646 |
| n315 | 0.0179270 | 0.2055643 | 0.7172150 | 0.5224277 | 0.7396370 | 0.9000840 | 1.503114e−01 | 0.387700119 |
| n316 | 13.2094277 | 14.6894373 | 16.1694469 | 16.1210393 | 17.5768451 | 18.9842434 | 8.338881e+00 | 2.887712148 |
| n317 | 11.8365885 | 11.9751162 | 12.2007247 | 12.8140338 | 13.0396423 | 15.0180974 | 2.209995e+00 | 1.486605063 |
| n318 | 9.4190176 | 10.6757340 | 10.6757405 | 10.5686663 | 10.9944380 | 11.0784013 | 4.464330e−01 | 0.668156385 |
| n319 | 4.2830280 | 4.8529163 | 5.0614500 | 4.9094000 | 5.1179338 | 5.2316720 | 1.810431e−01 | 0.425491595 |
| n320 | 6.9250485 | 7.2321927 | 7.5393369 | 7.5393369 | 7.8464812 | 8.1536254 | 7.547006e−01 | 0.868735040 |
| n321 | 2.8291110 | 2.9538207 | 3.0785304 | 3.0785304 | 3.2032401 | 3.3279498 | 1.244201e−01 | 0.352732322 |
| n322 | 2.6272780 | 2.7459529 | 2.8646278 | 2.8646278 | 2.9833026 | 3.1019775 | 1.126698e−01 | 0.335663235 |
| n323 | 2.4000180 | 2.5057123 | 2.6114065 | 2.6114065 | 2.7171008 | 2.8227950 | 8.937020e−02 | 0.298948484 |
| n324 | 11.8826648 | 11.8826648 | 11.8826648 | 11.8826648 | 11.8826648 | 11.8826648 | NA | NA |
| n325 | 9.8622455 | 11.0682280 | 11.3986431 | 13.6433549 | 12.5133867 | 22.6332555 | 2.666495e+01 | 5.163811676 |
| n326 | 9.4964128 | 9.9266021 | 10.3567914 | 10.3567914 | 10.7869807 | 11.2171701 | 1.480503e+00 | 1.216759161 |
| n327 | 6.9919767 | 7.9814300 | 11.5091574 | 11.1793396 | 14.7070670 | 14.7070670 | 1.688323e+01 | 4.108920366 |
| n328 | 4.4424450 | 4.8402684 | 5.2070772 | 5.1885988 | 5.5842609 | 5.8470060 | 2.881811e−01 | 0.536825063 |
| n329 | 3.7009810 | 4.2364910 | 4.9976954 | 5.8075639 | 8.0513260 | 8.0513260 | 4.407696e+00 | 2.099451436 |
| n330 | 4.0049850 | 4.4009303 | 4.7968755 | 4.7639842 | 5.1434838 | 5.4900920 | 5.521971e−01 | 0.743099644 |
| n331 | 2.3740030 | 2.9874095 | 3.3108200 | 3.1104665 | 3.3519825 | 3.4096585 | 1.441125e−01 | 0.379621513 |
| n332 | 2.4262040 | 2.5472593 | 2.6683145 | 2.6683145 | 2.7893698 | 2.9104250 | 1.172350e−01 | 0.342395953 |
| n333 | 3.0672930 | 3.3536390 | 3.4537530 | 3.6368540 | 3.6473070 | 4.6622780 | 3.724350e−01 | 0.610274547 |
| n334 | 3.3174200 | 3.4785386 | 3.6396573 | 3.6396573 | 3.8007759 | 3.9618945 | 2.076737e−01 | 0.455712289 |
| n335 | 2.9082070 | 3.0375515 | 3.1668960 | 3.1668960 | 3.2962405 | 3.4255850 | 1.338400e−01 | 0.365841492 |
| n336 | 6.4901043 | 6.7548642 | 7.0196240 | 7.0782358 | 7.3723016 | 7.7249792 | 3.838055e−01 | 0.619520358 |
| n337 | 5.1643530 | 5.8660845 | 6.2267450 | 6.0402515 | 6.4009120 | 6.5431630 | 3.739363e−01 | 0.611503321 |
| n338 | 5.6971505 | 6.2073208 | 6.4725678 | 6.3598579 | 6.6251049 | 6.7971455 | 2.246436e−01 | 0.473965868 |
| n339 | 3.3141840 | 3.7310860 | 3.7967050 | 4.8443528 | 5.0251008 | 8.2226890 | 4.382251e+00 | 2.093382637 |
| n340 | 6.5713000 | 7.8478863 | 7.9214353 | 7.7422748 | 7.9793078 | 8.2175098 | 3.468375e−01 | 0.588929123 |
| n341 | 4.9286330 | 5.6698280 | 5.6862690 | 5.6246557 | 5.8135810 | 5.9263970 | 1.276384e−01 | 0.357265112 |
| n342 | 4.5179220 | 5.0945168 | 5.3404645 | 5.1608469 | 5.4067947 | 5.4445368 | 1.880451e−01 | 0.433641688 |
| n343 | 3.9094410 | 4.3257480 | 4.5311558 | 4.4066016 | 4.6120094 | 4.6546540 | 1.162025e−01 | 0.340884914 |
| n344 | 8.5866236 | 9.4071402 | 10.2276568 | 10.0089100 | 10.7200532 | 11.2124495 | 1.759628e+00 | 1.326509731 |
| n345 | 6.9820981 | 7.6624323 | 8.3427665 | 8.4206529 | 9.1399303 | 9.9370941 | 2.187550e+00 | 1.479036866 |
| n346 | 6.2848971 | 7.2289350 | 7.4579823 | 9.0258784 | 8.2550814 | 15.5567620 | 1.212785e+01 | 3.482505886 |
| n347 | 5.7308092 | 6.2519884 | 6.3827302 | 6.3529669 | 6.5256355 | 6.8398533 | 1.390683e−01 | 0.372918563 |
| n348 | 5.0843193 | 5.7024969 | 6.0309493 | 6.0592220 | 6.3876744 | 7.0906700 | 6.819597e−01 | 0.825808506 |
| n349 | 4.1320500 | 4.7315119 | 4.9254013 | 4.7752658 | 4.9284171 | 5.0724380 | 1.165261e−01 | 0.341359125 |
| n350 | 0.5650320 | 1.4501485 | 1.9739585 | 2.3049306 | 2.7134388 | 4.7551030 | 2.619991e+00 | 1.618638543 |
| n351 | 3.1757560 | 3.1989679 | 3.2695798 | 3.3624886 | 3.3468445 | 3.9119860 | 7.780931e−02 | 0.278943194 |
| n352 | 0.1516230 | 0.2210645 | 0.2732460 | 0.5734643 | 0.4287145 | 2.2898230 | 5.914059e−01 | 0.769029169 |
| n353 | 3.7234140 | 3.9384905 | 4.0861402 | 4.0800156 | 4.1776262 | 4.4870037 | 6.913450e−02 | 0.262934398 |
| n354 | 1.7432180 | 1.7562284 | 1.8972510 | 2.0541300 | 2.2269493 | 2.7409905 | 1.593998e−01 | 0.399249020 |
| n355 | 1.7753940 | 1.9357835 | 2.0961730 | 2.1186286 | 2.2902458 | 2.4843187 | 1.260217e−01 | 0.354995402 |
| n356 | 4.9105590 | 5.7381080 | 5.8217245 | 5.8317691 | 5.9094050 | 6.7790490 | 4.402170e−01 | 0.663488486 |
| n357 | 3.3458490 | 3.5768375 | 3.6836130 | 3.8300510 | 3.8255345 | 4.9761510 | 2.884649e−01 | 0.537089290 |
| n358 | 17.0080395 | 17.6076681 | 27.0393487 | 26.8394725 | 36.2711531 | 36.2711531 | 1.187153e+02 | 10.895656643 |
| n359 | 14.2560300 | 14.3195990 | 14.3831680 | 14.7251200 | 14.9596650 | 15.5361620 | 4.973829e−01 | 0.705253754 |
| n360 | 14.6542715 | 16.2028393 | 17.5090652 | 20.4256522 | 22.2598771 | 31.3449732 | 4.749813e+01 | 6.891889052 |
| n361 | 12.7860291 | 14.3892912 | 16.7657896 | 18.9204081 | 22.7720291 | 28.5683972 | 4.619521e+01 | 6.796705464 |
| n363 | 11.6860470 | 11.6860470 | 11.6860470 | 11.6860470 | 11.6860470 | 11.6860470 | NA | NA |
| n364 | 8.4738420 | 9.7376363 | 10.3437614 | 13.5061004 | 19.4876312 | 19.4876312 | 3.027069e+01 | 5.501880488 |
| n365 | 8.1097906 | 8.4883897 | 8.6872947 | 8.7136202 | 8.9125252 | 9.3701008 | 2.691784e−01 | 0.518824026 |
| n366 | 6.9135516 | 7.0610176 | 11.4796762 | 11.4305208 | 15.8491793 | 15.8491793 | 2.603917e+01 | 5.102858743 |
| n368 | 4.9104490 | 4.9104490 | 4.9104490 | 4.9104490 | 4.9104490 | 4.9104490 | 0.000000e+00 | 0.000000000 |
| n369 | 4.7649100 | 5.0265268 | 5.2881435 | 5.2234535 | 5.4527252 | 5.6173069 | 1.847837e−01 | 0.429864757 |
| n370 | 1.5599040 | 1.7290031 | 1.9450100 | 1.9181350 | 2.1255028 | 2.2162180 | 7.167838e−02 | 0.267728175 |
| n371 | 0.2330240 | 0.3094468 | 0.5554813 | 0.5225778 | 0.5722479 | 0.9862080 | 7.756608e−02 | 0.278506871 |
| n372 | 0.2330240 | 0.2624585 | 0.3616790 | 0.3341822 | 0.3725960 | 0.4430910 | 7.110644e−03 | 0.084324634 |
| n373 | 0.1042900 | 0.1042900 | 0.3173370 | 0.3255905 | 0.5386375 | 0.5633980 | 6.548020e−02 | 0.255890997 |
| n374 | 6.0861970 | 6.0861970 | 6.0861970 | 6.0861970 | 6.0861970 | 6.0861970 | NA | NA |
| n375 | 3.0623990 | 3.0623990 | 3.0623990 | 3.0623990 | 3.0623990 | 3.0623990 | NA | NA |
| n377 | 4.2961310 | 4.6080905 | 4.8587307 | 4.8148238 | 5.0654640 | 5.2457030 | 1.671898e−01 | 0.408888464 |
| n385 | 3.6759030 | 3.7331053 | 3.7903075 | 3.7903075 | 3.8475098 | 3.9047120 | 2.617678e−02 | 0.161792395 |
| n386 | 1.7372710 | 1.7643053 | 1.7913395 | 1.7913395 | 1.8183738 | 1.8454080 | 5.846805e−03 | 0.076464406 |
| n390 | 1.7881500 | 1.8716250 | 1.8994520 | 1.8952043 | 1.9230313 | 1.9937630 | 7.070175e−03 | 0.084084335 |
| n391 | 2.9777210 | 3.1167336 | 3.5920471 | 3.5700190 | 4.0453324 | 4.1182607 | 3.401323e−01 | 0.583208651 |
| n392 | 1.6924020 | 1.7193015 | 1.7462010 | 1.7462010 | 1.7731005 | 1.8000000 | 5.788665e−03 | 0.076083275 |
| n393 | 2.2791340 | 2.3855339 | 2.7550765 | 2.7740718 | 3.1436144 | 3.3070000 | 2.509704e−01 | 0.500969421 |
| n394 | 1.3589040 | 1.4079248 | 1.4685920 | 1.4522520 | 1.5129193 | 1.5129200 | 5.619404e−03 | 0.074962685 |
| n397 | 5.1930090 | 5.4354398 | 5.5162505 | 6.1475023 | 6.2283130 | 8.3644990 | 2.207696e+00 | 1.485831922 |
| n401 | 3.0016160 | 3.1417438 | 3.8423025 | 3.8082835 | 4.5088423 | 4.5469130 | 6.845463e−01 | 0.827373135 |
| n402 | 2.4789150 | 2.5946415 | 3.0204825 | 3.0660718 | 3.4919128 | 3.7444070 | 3.696659e−01 | 0.608001550 |
| n403 | 3.1913680 | 3.3216100 | 3.3775203 | 3.3341071 | 3.3900174 | 3.3900200 | 9.194140e−03 | 0.095886076 |
| n405 | 5.1229480 | 5.3621088 | 5.4418295 | 5.3666435 | 5.4463643 | 5.4599670 | 2.646755e−02 | 0.162688497 |
| n406 | 2.0570810 | 2.1531103 | 2.1851225 | 2.1763393 | 2.2083515 | 2.2780310 | 8.239344e−03 | 0.090770834 |
| n409 | 11.4013560 | 11.4013560 | 11.4013560 | 11.4013560 | 11.4013560 | 11.4013560 | NA | NA |
| n410 | 3.5493080 | 3.6045403 | 3.6597725 | 3.6597725 | 3.7150048 | 3.7702370 | 2.440481e−02 | 0.156220394 |
| n411 | 11.2726590 | 11.5169389 | 12.6935734 | 12.8267197 | 14.0033542 | 14.6470729 | 2.721069e+00 | 1.649566310 |
| n412 | 10.7125645 | 10.7125645 | 10.7125645 | 10.7125645 | 10.7125645 | 10.7125645 | NA | NA |
| n413 | 9.7765326 | 9.7765326 | 9.7765326 | 9.7765326 | 9.7765326 | 9.7765326 | NA | NA |
| n414 | 6.3687046 | 6.6941536 | 7.0196025 | 6.9482830 | 7.2380723 | 7.4565420 | 2.996624e−01 | 0.547414276 |
| n415 | 5.2404429 | 6.2391916 | 7.2379402 | 6.6449744 | 7.3472401 | 7.4565400 | 1.491478e+00 | 1.221260767 |
| n416 | 2.1730943 | 2.2762288 | 2.3825185 | 2.3481408 | 2.4544305 | 2.4544320 | 1.821510e−02 | 0.134963321 |
| n417 | 1.2439873 | 1.3656746 | 1.4500030 | 1.4094408 | 1.4937693 | 1.4937700 | 1.386926e−02 | 0.117767813 |
| n418 | 0.1877050 | 0.1964680 | 0.1993890 | 0.2663501 | 0.2692711 | 0.4789175 | 2.011251e−02 | 0.141818580 |
| n419 | 0.1176100 | 0.1231008 | 0.1249310 | 0.1413810 | 0.1432113 | 0.1980520 | 1.439289e−03 | 0.037937966 |
| n420 | 8.7122036 | 8.7122036 | 8.7122036 | 8.7122036 | 8.7122036 | 8.7122036 | NA | NA |
| n421 | 5.8776403 | 5.8776403 | 5.8776403 | 5.8776403 | 5.8776403 | 5.8776403 | NA | NA |
| n422 | 5.4656660 | 5.7624820 | 5.8397770 | 5.8620532 | 6.1211700 | 6.1211710 | 7.545425e−02 | 0.274689377 |
| n423 | 21.1397190 | 21.1397190 | 21.1397190 | 21.1397190 | 21.1397190 | 21.1397190 | 0.000000e+00 | 0.000000000 |
| n424 | 5.1741564 | 6.6766753 | 8.1791943 | 7.2422469 | 8.2762922 | 8.3733900 | 3.217177e+00 | 1.793649008 |
| n425 | 3.5881220 | 3.5881220 | 3.5881220 | 3.5881220 | 3.5881220 | 3.5881220 | NA | NA |
| n426 | 3.3933560 | 3.5517718 | 4.6421313 | 4.6079464 | 5.6983059 | 5.7541670 | 1.648142e+00 | 1.283799875 |
| n427 | 3.0766020 | 3.4133725 | 3.7501430 | 3.5350250 | 3.7642365 | 3.7783300 | 1.578124e−01 | 0.397256046 |
| n428 | 0.8199778 | 1.3119990 | 1.3990550 | 1.2915484 | 1.4405700 | 1.4861400 | 7.360701e−02 | 0.271306121 |
| n429 | 1.4830000 | 1.5578833 | 1.6327665 | 1.6327665 | 1.7076498 | 1.7825330 | 4.486001e−02 | 0.211801815 |
| n431 | 9.8888710 | 10.1636431 | 10.4384153 | 10.4384153 | 10.7131874 | 10.9879595 | 6.039978e−01 | 0.777172954 |
| n432 | 9.0529051 | 9.0529051 | 9.0529051 | 9.0529051 | 9.0529051 | 9.0529051 | NA | NA |
| n436 | 7.4402441 | 7.4402441 | 7.4402441 | 7.4402441 | 7.4402441 | 7.4402441 | NA | NA |
| n437 | 6.3138615 | 6.6086190 | 7.1458488 | 7.0522485 | 7.5894783 | 7.6034350 | 4.173153e−01 | 0.645999480 |
| n438 | 5.0739840 | 5.1529423 | 5.2319005 | 5.2319005 | 5.3108588 | 5.3898170 | 4.987524e−02 | 0.223327656 |
| n439 | 5.6704717 | 5.6704717 | 5.6704717 | 5.6704717 | 5.6704717 | 5.6704717 | NA | NA |
| n441 | 2.2930062 | 2.2930062 | 2.2930062 | 2.2930062 | 2.2930062 | 2.2930062 | NA | NA |
| n443 | 6.6131064 | 6.6131064 | 6.6131064 | 6.6131064 | 6.6131064 | 6.6131064 | NA | NA |
| n445 | 4.7306712 | 4.7306712 | 4.7306712 | 4.7306712 | 4.7306712 | 4.7306712 | NA | NA |
| n449 | 4.4728941 | 4.4728941 | 4.4728941 | 4.4728941 | 4.4728941 | 4.4728941 | NA | NA |
| n450 | 8.3760986 | 8.3760986 | 8.3760986 | 8.3760986 | 8.3760986 | 8.3760986 | NA | NA |
| n452 | 6.1606798 | 6.4928131 | 6.8750201 | 6.8755188 | 7.2577259 | 7.5913553 | 3.902791e−01 | 0.624723209 |
| n453 | 4.6671898 | 5.0750423 | 5.3701131 | 5.3049572 | 5.7043797 | 5.7080610 | 1.964711e−01 | 0.443250629 |
| n454 | 4.2299997 | 4.5237758 | 4.6888012 | 4.6292461 | 4.7942715 | 4.9093823 | 8.465748e−02 | 0.290959590 |
| n455 | 4.2299996 | 4.2299996 | 4.2299996 | 4.2299996 | 4.2299996 | 4.2299996 | NA | NA |
| n458 | 2.5492680 | 2.5889380 | 2.6286080 | 2.6286080 | 2.6682780 | 2.7079480 | 1.258967e−02 | 0.112203704 |
| n459 | 1.4411250 | 1.4555925 | 1.4956215 | 1.5294450 | 1.5694740 | 1.6854120 | 1.229773e−02 | 0.110895117 |
| n460 | 1.4898300 | 1.5130140 | 1.5361980 | 1.5361980 | 1.5593820 | 1.5825660 | 4.299983e−03 | 0.065574254 |
| n461 | 1.4529230 | 1.4696923 | 1.4864615 | 1.4864615 | 1.5032308 | 1.5200000 | 2.249662e−03 | 0.047430602 |
| n462 | 2.0176235 | 2.0467134 | 2.2204040 | 2.2478119 | 2.4215025 | 2.5328160 | 6.316816e−02 | 0.251332772 |
| n463 | 1.3478240 | 1.5916198 | 1.7711450 | 1.7189703 | 1.8984955 | 1.9857670 | 7.789484e−02 | 0.279096464 |
| n464 | 3.1374925 | 3.2748119 | 3.6566140 | 3.6729721 | 4.0547743 | 4.2411680 | 2.786504e−01 | 0.527873449 |
| n465 | 0.9178650 | 1.0422353 | 1.1894025 | 1.1680995 | 1.3152668 | 1.3757280 | 4.299465e−02 | 0.207351517 |
| n466 | 1.0536064 | 1.3296465 | 2.2730385 | 2.9786262 | 4.6932630 | 5.7568870 | 4.408077e+00 | 2.099542018 |
| n467 | 13.9549165 | 14.5891893 | 14.8120768 | 14.6006545 | 14.8235420 | 14.8235480 | 1.854402e−01 | 0.430627691 |
| n469 | 10.5466010 | 10.5466010 | 10.5466010 | 10.5466010 | 10.5466010 | 10.5466010 | NA | NA |
| n470 | 3.2930800 | 3.7656010 | 3.9999930 | 3.9249248 | 4.1659550 | 4.3999950 | 1.785086e−01 | 0.422502784 |
| n471 | 8.5714722 | 10.4217920 | 11.0705030 | 10.5987030 | 11.3944980 | 11.5352500 | 1.468492e+00 | 1.211813466 |
| n472 | 7.6628417 | 7.6628417 | 7.6628417 | 7.6628417 | 7.6628417 | 7.6628417 | NA | NA |
| n473 | 5.5082202 | 5.5082202 | 5.5082202 | 5.5082202 | 5.5082202 | 5.5082202 | NA | NA |
| n474 | 7.6600520 | 7.6875704 | 7.7150889 | 7.7902950 | 7.8554164 | 7.9957440 | 3.241425e−02 | 0.180039568 |
| n475 | 10.4206740 | 10.7449920 | 11.0693100 | 10.8530998 | 11.0693128 | 11.0693155 | 1.402441e−01 | 0.374491757 |
| n476 | 8.6339120 | 8.9026210 | 9.1713300 | 8.9921925 | 9.1713328 | 9.1713355 | 9.627369e−02 | 0.310280015 |
| n477 | 2.3858440 | 2.3866428 | 2.4606295 | 2.4603638 | 2.5343505 | 2.5343520 | 7.299007e−03 | 0.085434224 |
| n478 | 4.9913250 | 5.1466675 | 5.3020100 | 5.1984493 | 5.3020115 | 5.3020130 | 3.217537e−02 | 0.179374934 |
| n482 | 8.0944590 | 8.3289754 | 8.5027238 | 8.4245525 | 8.5983008 | 8.5983033 | 5.654740e−02 | 0.237796971 |
| n483 | 6.3437690 | 6.4564730 | 6.6163405 | 6.5787726 | 6.7386401 | 6.7386405 | 3.784049e−02 | 0.194526320 |
| n484 | 4.0862400 | 4.2770025 | 4.3405905 | 4.3232058 | 4.3867938 | 4.5254020 | 3.254685e−02 | 0.180407455 |
| n486 | 1.9120460 | 1.9224365 | 1.9259000 | 1.9751678 | 1.9786313 | 2.1368250 | 1.165735e−02 | 0.107969200 |
| n487 | 3.4273970 | 3.5874028 | 3.6407390 | 3.6053143 | 3.6586505 | 3.7123820 | 1.520929e−02 | 0.123325959 |
| n488 | 3.3034594 | 5.1743365 | 7.4798020 | 6.6990396 | 8.3291610 | 9.1030210 | 6.008838e+00 | 2.451293182 |
| n489 | 3.3706610 | 3.3806953 | 3.9410660 | 4.0077178 | 4.5680885 | 4.7780780 | 5.429124e−01 | 0.736825863 |
